# A post-embryonic specific and ecologically selective network underlies metamorphic remodelling in teleosts

**DOI:** 10.64898/2026.07.29.741460

**Authors:** Agneesh Barua, Giulia Campli, Marcela Herrera, Saori Miura, Yo Y. Yamasaki, Ken Maeda, Yann Gibert, Vincent Laudet, Marc Robinson-Rechavi

## Abstract

Evolutionary developmental biology has primarily focused on embryogenesis. Yet, much of the diversity in animal form and function emerges during post-embryonic developmental transitions such as metamorphosis. Whether these transitions share a conserved genetic architecture, and how that architecture evolves remains unclear. In this study we use a comparative framework and multiple data modalities to characterise post-embryonic development in five species of teleost fish. Despite 200 million years of divergence, we uncover a post-embryonic developmental gene network conserved in all five species. This network comprises core cellular process modules shared with embryogenesis, and developmental remodelling modules unique to post-embryonic development. Both sets have highly conserved expression and tissue specificity patterns across several teleost species and the outgroup spotted gar. The network is broadly responsive to thyroid hormone treatments, yet its response to ecological variation is focussed and selective. Although largely conserved at the coding sequence level, the network has undergone significant turnover in gene family copy number following the teleost-specific whole-genome duplication. Across teleosts, the copy number variation is associated with macroevolutionary variation in a number of ecological and morphological traits such as trophic level and swimming performance. Multi-species tissue expression, and zebrafish perturbation phenotypes provide support for these associations. Our results reveal an ancient, endocrine controlled network underlying post-embryonic development in teleosts. It suggests a conserved, yet evolvable genetic architecture can be redeployed, refined, and repurposed to generate diversity in form and function.

**Sentence summary:** Teleost post-embryonic remodelling is carried out by a conserved yet evolvable gene regulatory network.

## Introduction

Early comparative studies described striking morphological similarities among embryos of different animals, showing that aspects of development are conserved across species (*1*, *2*). These ideas were later extended to the molecular genetic level through concepts such as the phylotypic stage (*3*, *4*), which has been demonstrated across a wide range of animal lineages (*5*). However, embryogenesis captures only part of the developmental basis of phenotypic evolution. In many animals, development continues well beyond the embryo, and post-embryonic development can involve extensive remodeling of morphology, physiology, and behavior (*6*). This is most evident in organisms with complex life cycles such as metamorphosis. During metamorphosis, larval phenotypes transform into juvenile or adult forms through coordinated, stage-specific gene regulatory programs (*7*). In anurans, thyroid-hormone-dependent transcriptional cascades drive the reabsorption of the tail, the growth of the limbs, and reorganisation of intestine, skin, and nervous system during the tadpole-to-frog transition (*8*). In insects, juvenile hormone and ecdysteroid signaling regulate the timing and execution of larval-to-adult transformation, underpinning the emergence of adult structures such as wings and reproductive morphology (*9*). In teleost fishes, metamorphosis similarly involves major reorganisation of body shape, pigmentation, feeding structures, sensory systems, and behavior, under the strong endocrine control of thyroid hormone (*10*).

A major discovery from comparative embryology was the spatial and temporal collinearity of *Hox* genes (*4*), a family of highly conserved genes regulating body patterning, whose copy number evolution can profoundly alter morphology (*11*). This led to the ‘genetic toolkit’ concept, which proposes that much of animal diversity emerged through modifications of conserved developmental regulators and signaling pathways, such as *Hox*, *Pax6*, *Wnt*, *Hedgehog* and *Notch* genes, rather than through the constant origin of entirely new developmental genes (*12*, *13*). For this reason, identifying conserved genes that regulate development has remained a major and continuing focus of evo-devo research. However, just as changes in embryonic patterning genes such as *Hox* can alter body organisation, changes in genes acting during post-embryonic development can also shape the final adult phenotype. This has evolutionary consequences because it is the reproductively mature adult that transmits genes to the next generation, and thus developmental processes occurring after embryogenesis can have major effects on the direction and outcome of evolution (*14*).

Teleost fish provide an especially powerful system to understand how post-embryonic developmental trajectories and metamorphic strategies are linked to ecology and life history (*7*, *10*, *15*). Across teleosts, metamorphosis underlies transitions between pelagic and benthic habitats, open ocean environments and coral reefs, freshwater and marine ecosystems in amphidromous, catadromous, and anadromous species (*16*), or dramatic shifts in diet, locomotion, pigmentation, and body organization (*17*). These transitions range from relatively subtle remodeling events to some of the most spectacular metamorphoses observed among vertebrates, such as the asymmetric body plan of flatfishes (*18*) the smoltification of salmonids (*19*), the larval-to-reef transition of coral reef fishes (*20*, *21*), or the transformation of leptocephalus larvae into juvenile eels (*22*). This striking diversity in form, function, and life history makes teleosts ideal for studying both the conserved and divergent molecular mechanisms that underlie vertebrate post-embryonic development and metamorphosis. In this study we construct developmental co-expression networks to classify development into modules of genes with coordinated expression across the post-embryonic transition of five teleosts. These modules reduce the complexity of development into biologically interpretable units and offer a tractable basis for comparing development across species, even when precise developmental stages are not perfectly conserved (*23*, *24*). The ability of network-based approaches to uncover the genetic basis of complex traits, including venom systems in amniotes (*25*), eyespot patterning in butterflies (*26*), and behavioral evolution in songbirds (*27*), demonstrates their broader potential for developmental biology.

We identify orthologous genes with conserved interaction structure across five fish species spanning ca. 200 million years of divergence, and uncover a conserved post-embryonic developmental network. This conserved co-expression network is largely unique to post-embryonic development. Parts of it are more responsive to ecological variation than others, suggesting a degree of ecological fine-tuning. Evolutionary variation within this network is also associated with multiple teleost phenotypes. Similar to how conserved developmental regulators have been central to understanding embryogenesis, our findings provide a powerful framework for understanding how vertebrate phenotypic diversity can evolve through changes in conserved post-embryonic developmental programmes.

## Results

### We use developmental and ecologically diverse species to identify deeply conserved features

In this study we characterise post-embryonic development in five species of teleosts from different families; clownfish (*Ampriprion ocellaris,* Pomacentridae), grouper (*Epinephelus malabaricus,* Serrinidae), manini (*Acanthurus triostegus,* Acanthuridae), zebrafish (*Danio rerio,* Danionidae), and goby (*Rhinogobius sp BB*, Oxudercidae) (Fig 1A). These species cover a wide range of ecological niches, life-history strategies, and developmental trajectories, allowing us to identify conserved molecular programs across multiple contexts. Clownfish have a biphasic life history during which the larvae are free swimming in the open ocean, then following metamorphosis juveniles colonise coral reefs and establish their iconic symbiosis with sea anemone (*15*). The clownfish specimens in our study were reared in animal husbandry and seven developmental stages representing complete larval development were sampled (*28*) (Fig 1A). There is a distinct peak of thyroid hormone expression at around the fourth developmental stage marking the onset of metamorphosis (*28*) (Fig 1A). Groupers have a similar transition from a free swimming larval stage to a reef-associated juvenile, but unlike the clownfish they represent one of the top predators of the reef food web (*29*). The grouper samples used in this study were raised in aquaculture and eight developmental stages representing complete larval development were sampled (*21*). They have two distinct thyroid hormone peaks, one at around three days post hatching (S2 in our study), and another around thirty four days (S6) post hatching (*21*) (Fig 1A). Manini are marine species that shift from planktonic feeding to benthic herbivores and typically inhabit highly heterogeneous coastal habitats (*20*). They have a large thyroid hormone peak while the fish is still in the open ocean, which is followed by the onset of reef recruitment (*20*). Unlike the clownfish and grouper, the manini samples used in this study were limited to the four larval stages at the onset of reef recruitment. Therefore, in our samples, we capture the late decline thyroid hormone peak, with the highest estimated concentration at the first sampled stage (*30*). These fish were wild caught and reared in aquaria (*30*). Many freshwater goby species are amphidromous, exhibiting a biphasic life history where they spend their larval stage in the sea before migrating upstream into freshwater streams and living a benthic life. However, *Rhinogobius sp*. *BB* spends its entire life in the upper reaches of the streams, and has evolved a large egg size in response to delayed hatching (*31*). For goby, we sampled seven stages from wild-caught specimens representing complete larval development. We could not estimate a clear thyroid hormone peak for goby, instead thyroid hormone levels increased steadily towards the end of development (*32*). Finally, we sampled complete larval development of the zebrafish lab strain AB, one of the most commonly used and well characterised zebrafish lines used in research (*33*, *34*). The thyroid hormone peak in zebrafish is estimated around three weeks post hatching (*35*), which in our samples was around the fifth sampled stage. These five species shared a common ancestor around 200 million years ago (Fig 1B and Fig S1)(*36*).

**Fig 1:**
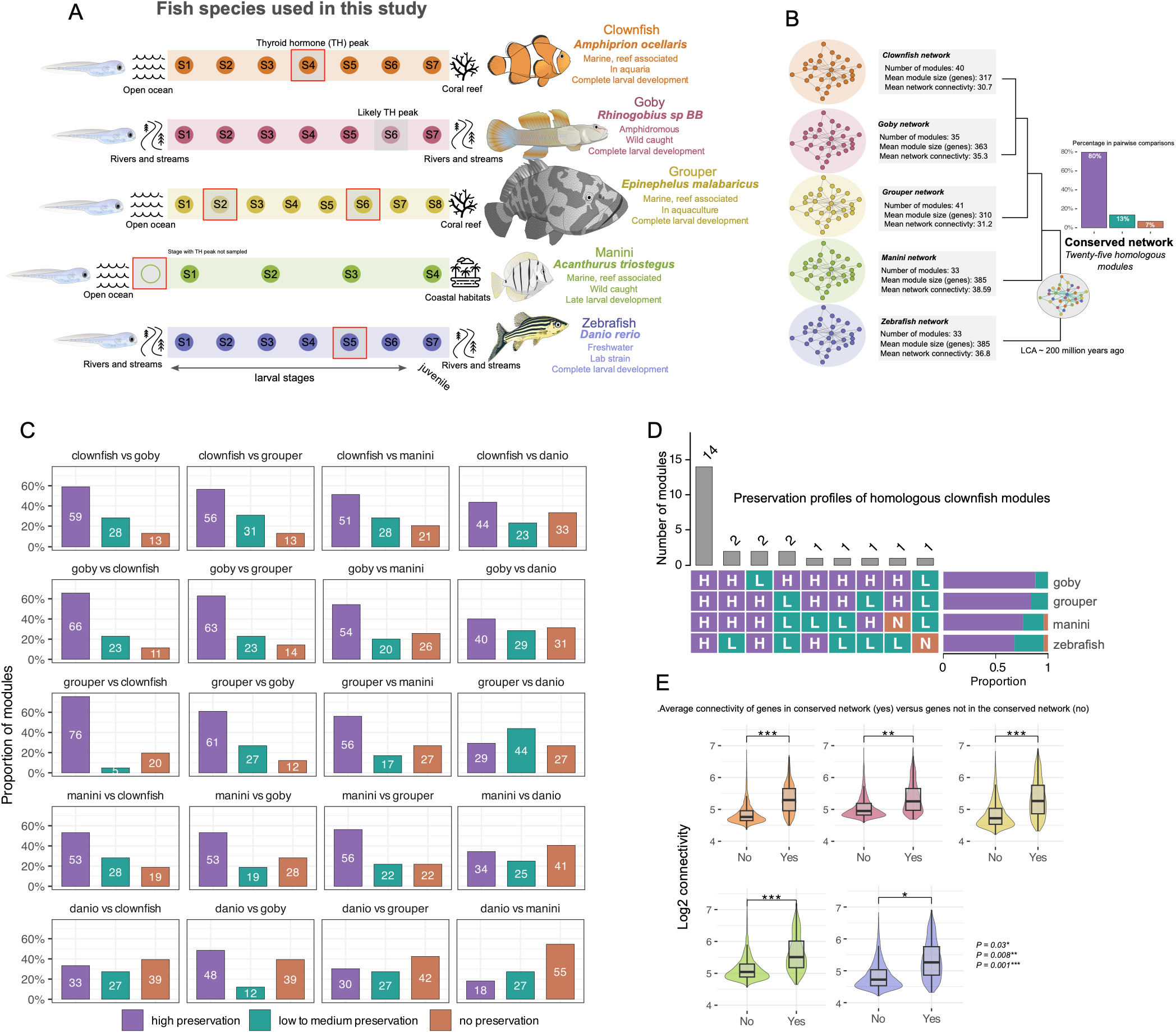
Conserved co-expression networks in teleosts and their network properties. (A) Diagram showing species used in this study along with their ecology, sample source, and sampling breadth. Diagrams are not to scale. The red boxes denote stages with measured thyroid hormone (TH) peaks. For goby the TH peak is an estimate while for manini, the stage with the TH peak was not sampled. The larvae and zebrafish diagram CC-BY Agneesh Barua, the others CC-BY Stefano Vianello. (B) The networks represent the individual species co-expression networks. The network diagrams are schematics and not the empirical networks. The networks are arranged according to the phylogenetic relationship of their respective species. The conserved post-embryonic developmental network is displayed at the root of the tree. The bar plot shows that 80% of the homologous modules in the conserved network are highly preserved among pairwise comparisons within individual species networks. (C) Module preservation between individual species networks. Module preservation estimates the degree to which interactions of orthologs in a reference network (first species name in panel), is conserved in a test network (second species name in panel). For example, in the first top left panel 59% of the modules in the clownfish network showed high preservation in the goby network, while 28% showed low preservation, and 13% showed no preservation. (D) The upset plot shows module preservation categories of the estimated homologous modules with clownfish modules as the reference. The tiles show each of the preservation profiles in the data with the corresponding bar plots showing their frequency. The bar plots on the right shows the proportion of homologous modules of each species belonging to each preservation category. (E) In each species, the genes making up the conserved network have an overall higher connectivity compared to genes not present in the conserved network.

### A conserved regulatory architecture underlies teleost post-embryonic development

Despite the differences in developmental strategies, timing, and ecology, we uncovered a conserved regulatory architecture across the five teleost species. This was based on 12,700 ortholog groups (see methods), which were then used to build a gene expression matrix for each species. This matrix was thus composed of the same ortholog groups in all five species. For orthologous groups containing multiple paralogs within a species, we selected the copy that had the highest expression across the developmental series, thereby capturing the major transcriptional contribution of each ortholog group (Fig S2). We then constructed a developmental network independently for each species (Fig 1B). This species-specific approach allowed us to circumvent the need for synchronous developmental stages. Instead, it enabled us to focus on identifying orthologous genes that retained coordinated expression across the broader larva-to-juvenile transition. The resulting networks contained between 33 and 41 gene modules, with the largest number recovered in grouper (Fig 1B). We confirmed that our network specifically groups genes due to their coordinated expression across development by comparing our network with one obtained from mean conserved permuted gene expression matrices (see methods).

A substantial proportion of the modules shared similar network properties across species. We quantified similarities in network properties using pairwise comparisons of the module preservation for all modules across all five species (*37*). Across species comparisons, most modules showed high preservation (Fig 1C). Clownfish and grouper showed the highest pairwise preservation, while zebrafish comparisons showed the lowest. Preservation of network structure does not necessarily mean that modules contain the same genes. Therefore, to identify modules with overlapping sets of genes we quantified module-membership and obtained module sets with a high degree of gene overlap across species. This analysis identified 25 cross-species homologous module sets. A large majority (80%) of the homologous module sets were preserved across pairwise comparisons demonstrating agreement between conservation of network structure and of gene composition (Fig 1B). For example, 14 out of 25 of the homologous modules in clownfish were highly preserved in the other four species, with a decrease in preservation across species following phylogenetic distance (Fig 1D). These 25 homologous module sets comprised a set of 1196 ortholog groups, together making up a conserved post-embryonic transcriptomic network. Orthologous groups belonging to this conserved network also had greater network connectivity than those outside the conserved network (two-sided Wilcoxon rank-sum test, Benjamini–Hochberg–adjusted P<0.05; Fig 1E). This indicated that they form central components of the shared transcriptomic architecture rather than a collection of weakly associated genes.

Together, these results reveal conserved transcriptomic components of post-embryonic development across five fish species. They provide evidence that post-embryonic development in teleosts can be characterised by conserved underlying transcriptomic network architectures.

### The conserved network is partitioned into core and remodelling processes

Gene Ontology (GO) enrichment (hypergeometric test, Benjamini-Hochberg adjusted P < 0.05) revealed that the 25 homologous module sets making up the conserved network were organised into two broad functional partitions (Fig 2A; FigS3; table S1). The first was the core cellular process partition, with processes such as cell cycle, RNA processing, and translation. The second was a remodelling partition comprising processes more directly associated with developmental remodelling and physiological maturation, such as nervous-system development, muscle-system development, eye development, and extracellular-matrix and cartilage formation. These modules were also expressed in similar tissues across multiple species of teleosts, as well as in the spotted gar (*Lepisosteus oculatus*) (Fig S4). Thereby, representing a highly conserved transcriptomic architecture.

**Fig 2:**
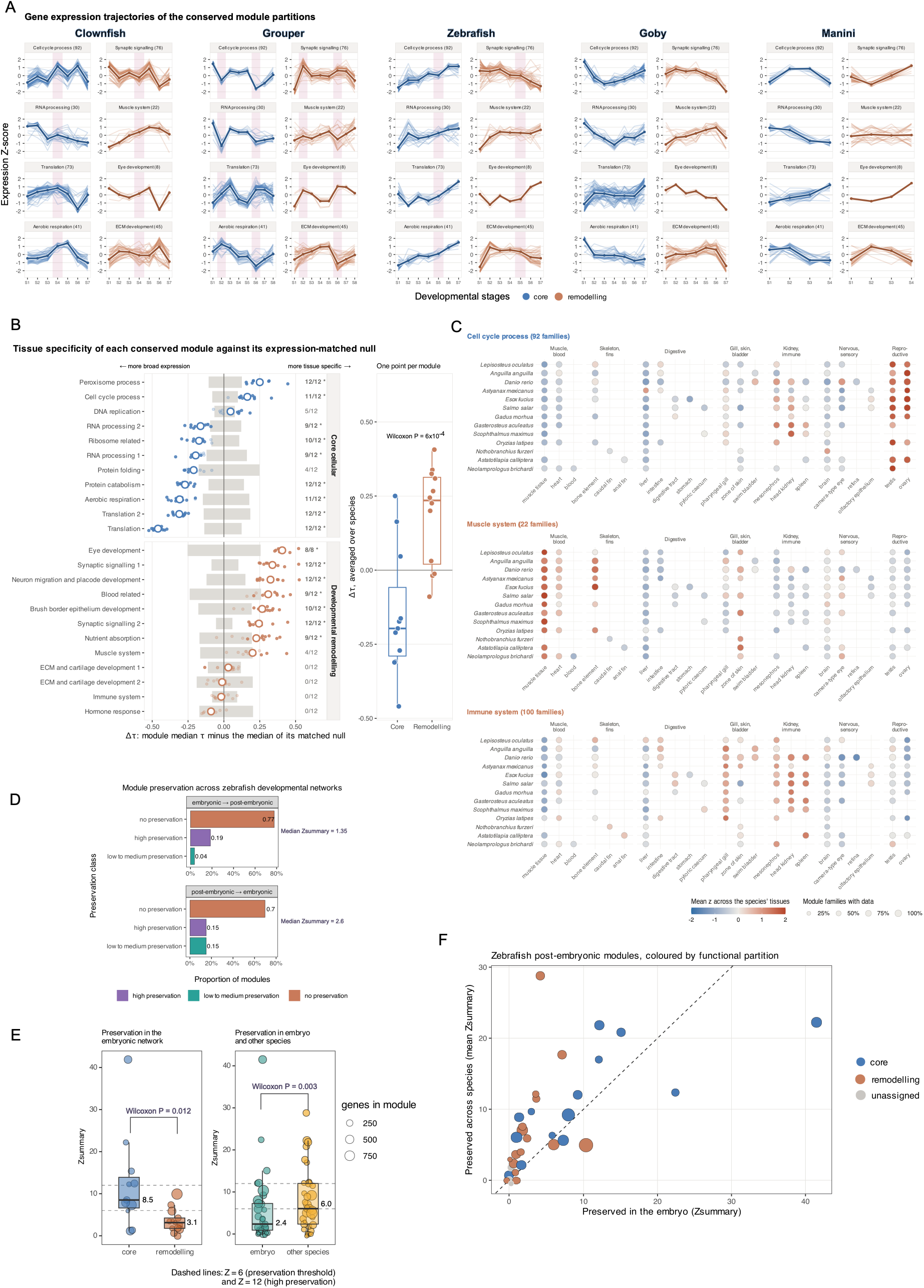
The conserved network is made of two highly conserved functional partitions and is specific to post-embryonic development. (A) Expression trajectories of ortholog groups making up the conserved module sets. For brevity only eight modules are shown. The blue panels are the core cellular partition and the brown panels are the remodelling partition. The red vertical bars denote peaks of thyroid hormone levels. There was an overall difference in gene expression trajectories of modules from the core cellular and remodelling partitions. (B) We calculated tissue specificity (tau) for each module and compared it against an expression matched genomic background. We report △tau which is the difference in median module tau minus the median background tau. The grey bars represent the 95% median intervals of the null centered around zero and averaged over all species. The large circle is the mean △tau over species. Small solid points are the △tau estimate in each species that lie outside the null distribution while the transparent points are △tau estimates that do not. The fractions are the number of species where the △tau estimate lies outside the null. The asterisks indicate modules where the difference from null occurs in at least half the species used in the comparison. The boxplots shows the difference in mean △tau between the two partitions. (C) Plots showing the combined expression of the modules across tissues in several species of teleosts and gar. The expression levels are plotted as z-scores computed across available tissues in each species. The numbers in brackets are the number of ortholog groups in each module. The size of the circles denote the percentage of the module with data. (D) Module preservation results between embryonic development network and post-embryonic network in zebrafish. (E) Network preservation: Left, significantly higher preservation of the conserved network core partition in embryogenesis. Right, significantly higher preservation of zebrafish post-embryonic modules in post-embryonic networks across species than in embryonic development within species. (F) Conserved network modules above the dashed line are better preserved across species, modules below the dashed line are better preserved in the same-species embryogenesis network.

These functional partitions differed in their expression dynamics across post-embryonic development. In clownfish, expression of genes within the core cellular partition generally decreased from early to late development (Fig 2A). By contrast, modules from the remodelling partition peaked either just after the peak in thyroid hormone levels, at the onset of metamorphosis, or during the later stages of metamorphosis (FigS3) Distinct trajectories were also observed in grouper. The expression of genes within the core cellular modules generally exhibited plateau-like trajectories between the two thyroid hormone peaks (Fig 2B; FigS3). On the other hand, genes within modules associated with eye development and synaptic signalling showed increased expression around both peaks. Some core programmes such as translation also increased around the second peak, suggesting redeployment of cellular machinery during later developmental remodelling.

The grouper trajectories also captured a lineage-specific morphological transformation. Pelvic floating spines begin to elongate after the first thyroid hormone peak and subsequently regress around the second peak (*21*). Mirroring these phenotypic changes, expression of the module involved in extracellular-matrix formation increased after the first peak and decreased during the second (Fig 2A). This pattern connects the activity of a conserved developmental module to a visible and species-specific remodelling change.

The aerobic respiration module captured a well described metabolic transition during metamorphosis. In clownfish, its expression increased after the thyroid hormone peak, around the onset of metamorphosis, consistent with the thyroid hormone–associated transition toward oxidative and lipid-based metabolism and the accompanying increase in mitochondrial gene expression (*28*). In grouper, the aerobic metabolic programme was most active earlier in larval development, around the first thyroid hormone peak, and subsequently decreased as metamorphosis progressed. This was consistent with observations of increased aerobic metabolic activity during the early stages of larval development with subsequent decrease as metamorphosis progressed (*21*). These observations show that conserved functional programmes can be deployed at different developmental times and with different temporal profiles, indicating heterochronic modification of a shared co-expression architecture.

In zebrafish, which does not undergo the extensive metamorphic transformation observed in clownfish and grouper (*33*), the distinction between the two functional partitions was less pronounced (Fig 2A). Module trajectories were either oscillatory, gradually increased over development, or showed comparatively subtle changes associated with thyroid hormone. Goby also showed different trajectories between core cellular and remodelling partitions. However, because thyroid hormone concentrations could not be reliably quantified, the endocrine signal associated with these changes could not be determined. In manini, despite sampling being restricted to late larval development, we observed differences in cell-cycle and RNA processing modules which decreased toward the end of the developmental series, whereas modules involved in eye development, synaptic signalling, neuronal migration, and nutrient absorption increased during reef recruitment (Fig 2A and Fig S3) Thus, the larva-to-juvenile transition involves a temporal progression from broad changes in core cellular processes to the deployment of distinct tissue-remodelling programmes.

To estimate how well the conserved network captures the functions of different biological systems, we calculated tissue specificity of its orthologs across adult tissues in several teleosts species and spotted gar. Genes of the core partition were, on average, more broadly expressed in the adult tissues than other expression matched genes from the same species, whereas genes of the remodelling partition were more tissue specific than the expression matched genomic background (Fig2B; expression matched permutation test; Benjamini-Hochberg-adjusted P < 0.05). Tissue specificity was also lower in the core cellular partition than in the remodelling partition (two-sided Wilcoxon rank sum test; P = 6×10^-4^) (Fig 2B). Within the core partition, the DNA replication, cell cycle, and peroxisome processes modules showed higher tissue specific than other modules. This was due to the high expression of the DNA replication and cell cycle modules in the gonads, and peroxisome module in the brain (Fig 2C, FigS4) Conversely, the muscle system, extracellular matrix remodelling, immune system, and hormone response modules showed relatively broader tissue expression than other remodelling partition modules. These patterns reflected their expression across tissues with related functions. For example, the muscle system module genes were expressed across muscle tissue, heart, bone, and skin, while the immune system module genes were expressed across the pharyngeal gills, mesonephros, head kidney, and spleen for the immune system modules (Fig 2C).

Together, these results revealed functional organisation underlying teleost post-embryonic development and the extent of its conservation. The functional organisation is partitioned into a core cellular component and a remodelling component. However, the timing and deployment of these partitions can vary among species. In clownfish and grouper, where hormone dynamics were sufficiently resolved, many of these changes were temporally associated with thyroid hormone peaks. Additionally, the orthologous genes of the conserved network have similar tissue expression patterns across multiple teleost species and the non-teleost spotted gar, along with similar levels of tissue specificity. Thus, the conserved network represents a highly conserved transcriptomic architecture that can undergo heterochronic shifts, and likely persisted since the common ancestor of teleosts.

### The remodelling partition is specific to post-embryonic development

Since the conserved network comprises a number of core cellular processes, the modules we observed could be a general property of highly conserved transcriptomes rather than a feature specific to post-embryonic development. To test this, we constructed a zebrafish embryonic co-expression network from a published developmental time-series (*38*), using the same orthologous groups, the exact same representative gene per group, and identical normalisation and network parameters applied to the post-embryonic networks. The majority of modules were not developmentally preserved between the two transitions. Of the 27 embryonic modules, 77% showed no preservation in the post-embryonic network, while of the 33 post-embryonic modules around 70% showed no preservation in the embryonic network (Fig 2C). These patterns were stable across multiple soft-thresholding powers (Table S2).

The core partitions had significantly higher preservation (two sided Wilcoxon rank sum test, P = 0.012) in the embryonic network than the remodelling partition (Fig 2D). The modules that were preserved in both the networks belonged to the core partition and were involved in translation, ribosome biogenesis, RNA processing, and DNA replication (Fig S5). The conserved network therefore comprises a functional partition of core cellular processes shared with embryogenesis, while the remodelling partition is unique to post-embryonic development.

Using the same set of orthologous groups as the above analysis, we recomputed pairwise module preservations against post-embryonic developmental networks of all five species and zebrafish embryogenesis. The zebrafish post-embryonic modules were significantly more preserved with post-embryonic network modules from the other species, than with the same-species embryogenesis network modules (two sided Wilcoxon rank sum test, P = 0.003) (Fig 2D). Assigning the dominant functional partitions to the modules showed that a majority of the zebrafish remodelling modules had higher preservation in post-embryonic development across species than between embryogenesis and post-embryonic development within zebrafish (Fig 2E).

Our results show that developmental gene modules of post-embryonic development are largely distinct from zebrafish embryogenesis. Furthermore, we show that post-embryonic modules are better preserved across 200 million years of species divergence than between embryonic and post-embryonic transitions within the same species. Together these results indicate a high degree of specificity of gene activity during post-embryonic development and their deep evolutionary conservation.

### The conserved network is highly responsive to thyroid hormone

We experimentally tested whether thyroid hormone can influence the conserved network by exposing clownfish to a treatment of exogenous triiodothyronine (T3), and to an inhibitor treatment of goitrogens called MPI (methimazole, potassium perchlorate, iopanoic acid). We compared gene expression differences between these treatments, with DMSO-treatment controls. Following T3 exposure, 66.8% of genes belonging to the conserved network genes were differentially expressed (n = 651, FDR < 0.05)(Fig 3A). Since the conserved network represents genomically conserved sets of orthologous genes, we compared the differential gene expression (DGE) estimates against two backgrounds: the ortholog groups used to construct the network but not in the conserved network, and a transcriptome background of all genes. The conserved network genes also have higher connectivity, which can covary with expression level. To account for this we created expression-matched random gene sets and counted the number of DEGs present in each set. We repeated this 10,000 times to get a null distribution of expression matched DEG rate. The observed DGE rate of 66.8% falls entirely above the expression matched null distribution for both the ortholog group and transcriptome backgrounds (Fig 3A). To complement this test and account for potential differences in level of dispersion between conserved network genes and non-network genes, we used a logistic regression adjusting for average expression with a natural spline and for gene-wise dispersion. The model gave an adjusted odds ratio of 1.77 (95% CI 1.54 - 2.05, P = 2.7 x10^-15^) against the ortholog groups and 1.69 (95% CI 2.47 - 1.94; P = 2×10^-13^) against the transcriptome background (Fig 3B). Thus, the conserved network is strongly enriched for differentially expressed genes following T3 treatment, even after accounting for genomic conservation and average expression level.

**Fig 3:**
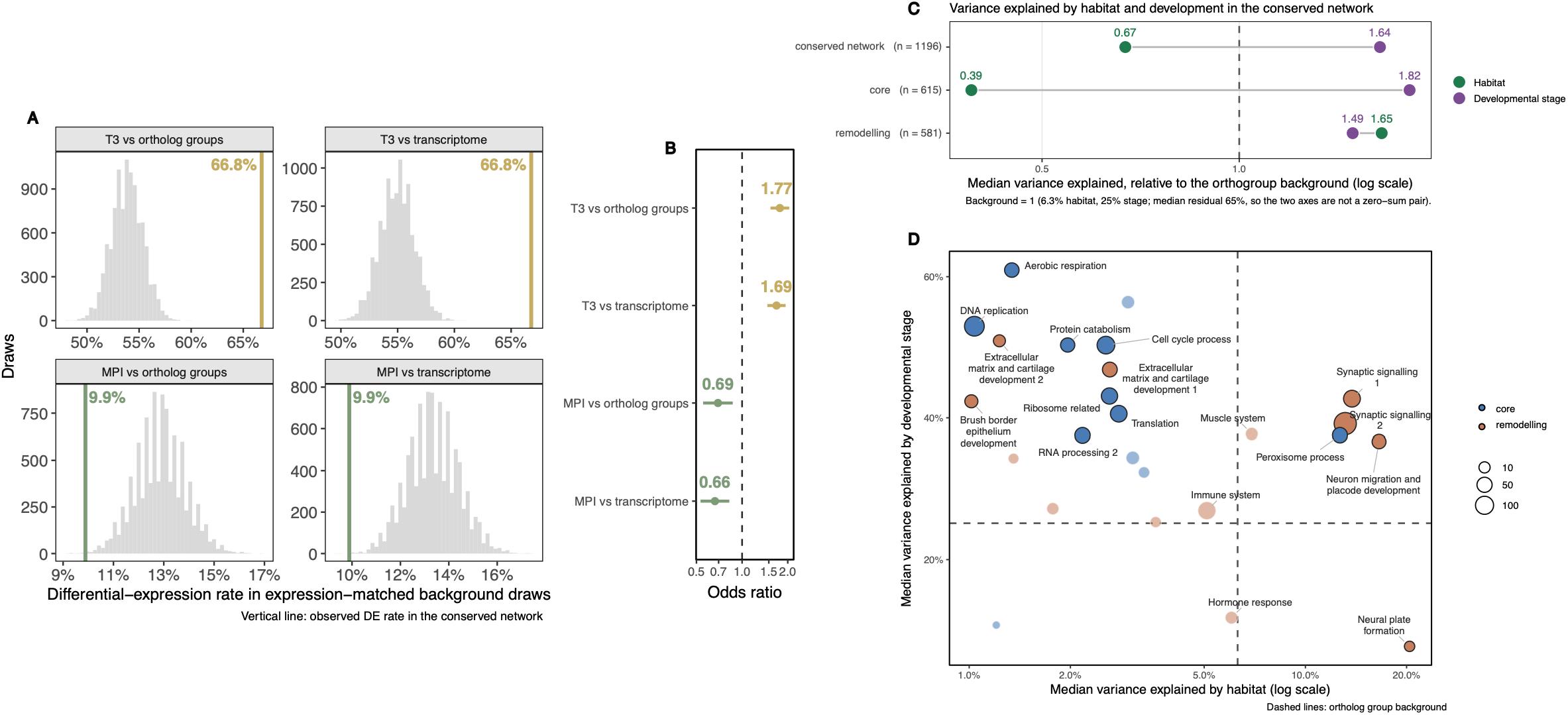
The conserved network is highly responsive to thyroid hormone (TH) and selectively responsive to habitat. **(A)** Plots showing the observed differential gene expression (DGE) rate of the conserved network against an expression matched null distribution using two background sets. For the T3 treatment the observed rate is well above the null rate for both backgrounds demonstrating that the conserved network is particularly responsive to TH induction. For the MPI treatment the observed rate is slightly below the null. (B) Odd ratios of a logistic regression of differential-expression status adjusted for average expression level. Against both backgrounds the conserved network was enriched for DGEs in the T3 treatment. Contrarily, the conserved network was depleted for DGEs in the MPI treatment. (C) Variance explained by habitat and development varied between partitions of the conserved network. Median ortholog group habitat variance: 0.063, conserved network habitat variance: 0.25; median ortholog group developmental variance: 0.25, median conserved network developmental variance: 0.41, (shuffled label permutation test, 10,000 iterations; P = 1×10^-4^); median core habitat variance: 0.024; median core developmental variance: 0.46; (shuffled label permutation test; P = 1×10^-4^); median remodelling habitat variance: 0.1; median core developmental variance: 0.37; (shuffled label permutation test P = 1×10^-4^). (D) Development explains much of the variance in gene expression of the core partition than habitat (left top corner). Habitat explains expression variance predominantly in neural modules of the remodelling partition and peroxisome process. Development also explains a substantial portion of variance in these modules suggesting that they respond to both developmental progression and habitat variation.

For the MPI treatment, only 9.9% of the genes of the conserved network were differentially expressed (n = 96, FDR < 0.05). The observed DGE rate was marginally lower than the expression-matched null distribution (Fig 3A). The logistic regression confirmed a significant depletion against both the ortholog group (odds ratio = 0.69; 95% CI 0.55 - 0.87; P = 1.4×10^-3^) and the transcriptome backgrounds (odds ratio = 0.66; 95% CI 0.53 - 0.82; P = 2.2×10^-4^) (Fig 3B). Across the ortholog group and transcriptome, the genes moved in opposite directions under the two treatments (Spearman rho =-0.33,-0.35 respectively; P= 2.2×10^-16^), however the conserved network showed a weaker directional relationship (Spearman rho =-0.14; P= 2.2×10^-16^) (Fig S6). The conserved network therefore responds asymmetrically to the treatments. It responds strongly to induction of TH using exogenous T3 but responds comparatively weaker to its suppression. This is consistent with the documented capacity of many fish (e.g. flatfish, fathead minnow, *A. ocellaris*) to compensate for the effects of goitrogenic compounds, which can only induce delay in metamorphosis without being able to completely block it (*39–42*).

Within the network, not all modules showed the same degree of responsiveness. The core partition genes had a stronger response to T3 induction than the remodelling genes (74.8 % versus 57.2% DEG, Fig S6; median |log₂FC| 0.539 versus 0.397; permutation test P = 1×10^-4^, Fig S6).

### The conserved network responds selectively to ecology

To test whether ecological context modulated the conserved network we used an *in-situ* rearing experiment where reef settled manini from a single site were reared in three distinct nursery habitats and sampled over a period of 8 days. The habitats, - mangrove, sand beach, and rock beach-in the island of Moorea in French Polynesia, were highly heterogeneous in both biotic and abiotic conditions (*30*).

We compared gene expression across fish from a time-series comprising stages S2, S3, and S4 reared in the three habitats. For each gene, we modelled expression as a function of developmental stage, or both habitat and developmental stage, and measured the gain in proportion of variance explained when including habitat in the model (see Methods). This was done to distinguish between the effects of development and habitat which cannot be estimated solely by single-stage pairwise comparisons between habitats. The two components separated expression variance differently across gene sets. Relative to the expression-matched ortholog group background genes, development explained more variation in the core partition as compared to habitat (Fig 3C). However, in the remodelling partition, both development and habitat explained more variance compared to the background. These broad differences were underlain by variation between individual modules (Fig 3D; Fig S6). Habitat predominantly explained variation in four nervous system related modules (remodeling partition) and the peroxisome module (core partition). Interestingly, these modules jointly vary across both development and habitat.

Together the results show that ecological context acts on the conserved network selectively. In manini, ecology selectively acts on the peroxisome module of the core partition, and on the nervous system modules of the remodelling partition. The majority of the core partition could be buffered from ecological effects through developmental canalisation, while the nervous system and peroxisome modules could be underlying habitat-driven phenotypic plasticity. Elevated levels of neurotransmitter γ-aminobutyric acid (GABA), and appetite regulatory profiles in the manini juveniles across habitat adds support to the latter (*30*, *43*).

### Genes of the conserved network are constrained in sequence but dynamic in copy number evolution

To determine how the conserved developmental network has evolved, we examined both coding-sequence evolution and gene-family turnover across 133 teleost species, using spotted gar as an outgroup. We compared the 1,196 ortholog groups belonging to the conserved network with an equally sized random set of ortholog groups outside the network. The genes belonging to the two sets did not differ significantly in gene length or ortholog copy number (bootstrapped two-sided Wilcoxon rank-sum tests, Benjamini–Hochberg–adjusted P > 0.05; Fig S7).

Overall dN/dS was similar between the conserved and null sets (Fig S8). Site and gene-level analyses identified slightly stronger purifying selection (χ² = 651.4, P < 2.2×10⁻¹⁶) together with a slight excess of genes showing episodic diversifying selection in the conserved network (two-sided Fisher’s exact test P = 0.011), but both effects were small (Cramér’s V = 0.019; odds ratio = 1.1).

We next used CAFE5 (*44*) to estimate gene copy expansions and contractions among the 1,196 ortholog groups across the teleost phylogeny and compared their frequencies with that of the null set. CAFE identified multiple instances of ortholog group expansion and contraction across the teleost phylogeny. We only report results for 1,104 ortholog groups as the analysis did not converge for the rest due to sparsity of the gene count matrix. Numerous expansions were inferred near the base of the teleost radiation, consistent with the retention of duplicated genes following the teleost-specific whole-genome duplication (*45*), followed by more lineage-specific expansions and contractions (Fig 4A). At the base of Anguilliformes (eels and tarpon), expansions involved ortholog groups associated with translation and synaptic signalling, whereas expansions at the base of Scombriformes (tunas and mackerels) were concentrated in immune-system, aerobic respiration, and synaptic-signalling modules (Fig 4B). By contrast, Siluriformes (catfishes) showed substantial contractions, particularly among ortholog groups associated with synaptic signalling. Contractions at the base of Syngnathiformes (seahorses and pipefish) involved ortholog groups belonging to immune-system and synaptic-signalling modules (Fig 4B). These changes show that lineages experienced copy number changes in both partitions of the conserved network. The complete list of high-turnover ortholog groups is provided in table S3.

**Fig 4:**
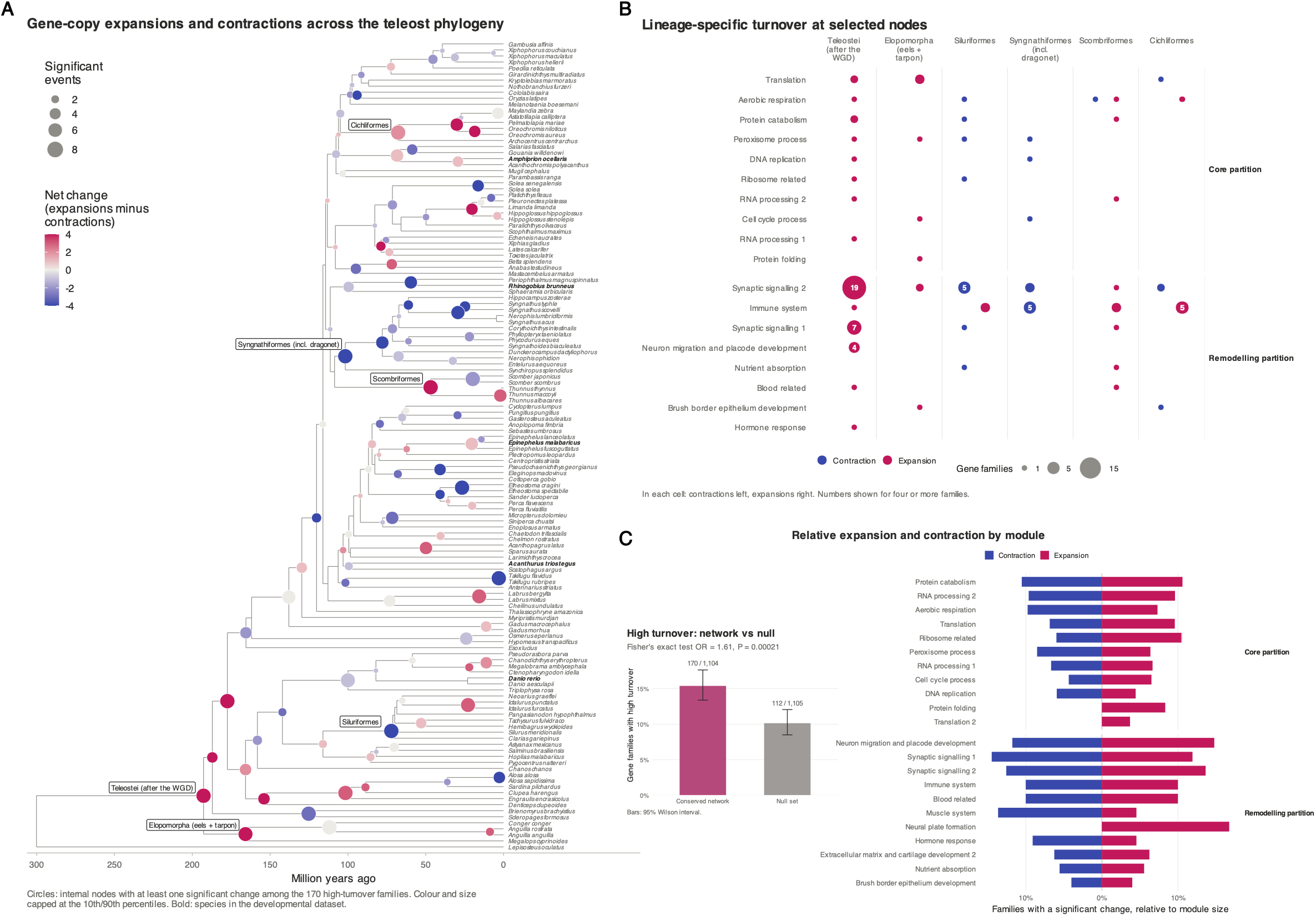
Evolutionary dynamics of gene copy evolution. (A) Expansion and contraction in gene copies of 170 ortholog groups of the conserved network mapped across the phylogeny of 133 teleost species with the spotted gar as an outgroup. The size of the circles at the nodes denotes the magnitude of gene copy number change, with the net direction (expansion or contraction) denoted by the color of the node. (B) Expansion and contraction of gene families from different modules at specific nodes in the teleost phylogeny. (C) The left plot shows the difference in proportion of high turnover genes comprising the conserved network compared to the random null set. The proportion of high turnover genes is greater in the conserved network. (Wilson interval: confidence interval for binomial proportions). The right plot shows the relative expansion and contraction of modules adjusted for its size.

Across the conserved network, 170 of 1,104 ortholog groups (15%) showed significant evidence of high turnover (Fig 4C). High-turnover groups were enriched in the conserved network than in the null set (two-sided Fisher’s exact test, odds ratio = 1.61, P=2×10−4). This enrichment was higher than the differences detected in the coding-sequence analyses. After accounting for differences in module size, ortholog groups associated with synaptic signalling, neuronal migration, and muscle-system development showed the greatest relative turnover (Fig 3B).

These results revealed two aspects of how the conserved network evolved. The protein-coding sequences of its genes have remained predominantly constrained. At the same time, the number of gene copies of the ortholog groups making up the conserved network have changed repeatedly across the teleost phylogeny. Such changes can provide a potential route through which a shared developmental system could be modified to generate lineage-specific developmental and phenotypic variation.

### Gene-family turnover in the conserved network is associated with phenotypic variation

We next asked whether gene turnover in the conserved developmental network was associated with phenotypic variation across teleosts. Using phylogenetic generalised linear mixed models (PGLMMs) we examined nine ecological, life-history, and morphological traits: maximum lifespan, age at maturity, average water temperature, trophic level, growth rate, body shape, demersal habitat, caudal-fin aspect ratio, and lower-jaw length (*46*)(Table S4). We summarised variation in the copy numbers of high-turnover ortholog groups using the first two axes of a phylogenetic principal components analysis (PCA) (*47*). Using the phylogenetic PCA scores captures the major axes of variation in gene copy number and prevents over parametrisation of the model. For associations whose 95% credible intervals excluded zero, we inspected both model convergence and examined the underlying data, which jointly helped to minimise the risk of spurious associations. We then evaluated the functional contexts using two independent resources: the Bgee database and the Zebrafish Information Network (ZFIN) phenotypes, to determine which tissues and processes were affected by experimental perturbation of genes in the associated modules (*48*, *49*). While these complementary datasets cannot establish that copy-number changes caused the associated phenotypes, they can determine whether the candidate genes function in relevant biological contexts. Applying these criteria identified several associations. For brevity we focus on one noteworthy case in the main text, and two additional cases in the supplementary material (Supplementary text 1). The results for all modules–trait associations and model diagnostics are provided in Fig S9 and the online supplementary material respectively.

### High turnover genes of the aerobic respiration module are associated with swimming performance and feeding ecology

Copy-number variation in high turn-over genes of the aerobic-respiration module was associated with trophic level and caudal-fin aspect ratio (Fig 5A). Trophic level was negatively associated with PC2 of the gene-copy phylogenetic PC axes (Fig 5A). Copy-number variation in ortholog groups containing *micos10*, a component of the mitochondrial contact site and cristae organising system (MICOS), and *atp5pd*, a subunit of mitochondrial ATP synthase peripheral stalk subunit d (ATPase d), contributed strongly to this axis. Species experiencing a loss of micos10 gene copy occupied a lower trophic level (Fig 5A). Zooplanktivorous species, including the European sardine (*Sardina pilchardus*) and stone moroko (*Pseudorasbora parva*), which experienced a loss in *micos10* gene copy, occupied lower trophic levels (Fig 5A, 5B). By contrast, apex predators such as tunas (*Thunnus thynnus* and *Thunnus albacares*) and swordfish (*Xiphias gladius*) which occupied higher trophic levels maintained the ancestral gene copy number. Caudal-fin aspect ratio is a common indicator of swimming performance (*46*). Species with higher aspect ratios tended to have lower scores along PC1, which primarily represented copy-number variation in ortholog groups containing ATP synthase membrane subunit genes (*atpmea* and *atpmeb*), and Ovarian Carcinoma Immunoreactive Antigen domain-containing protein gene (OCIAD; *ociad1*) and (Fig 5A). The OCIAD ortholog group showed extensive copy-number variation across teleosts, whereas the ATP synthase membrane subunit genes experienced losses in several lineages (Fig 5B)(Supplementary text 1).

**Fig 5:**
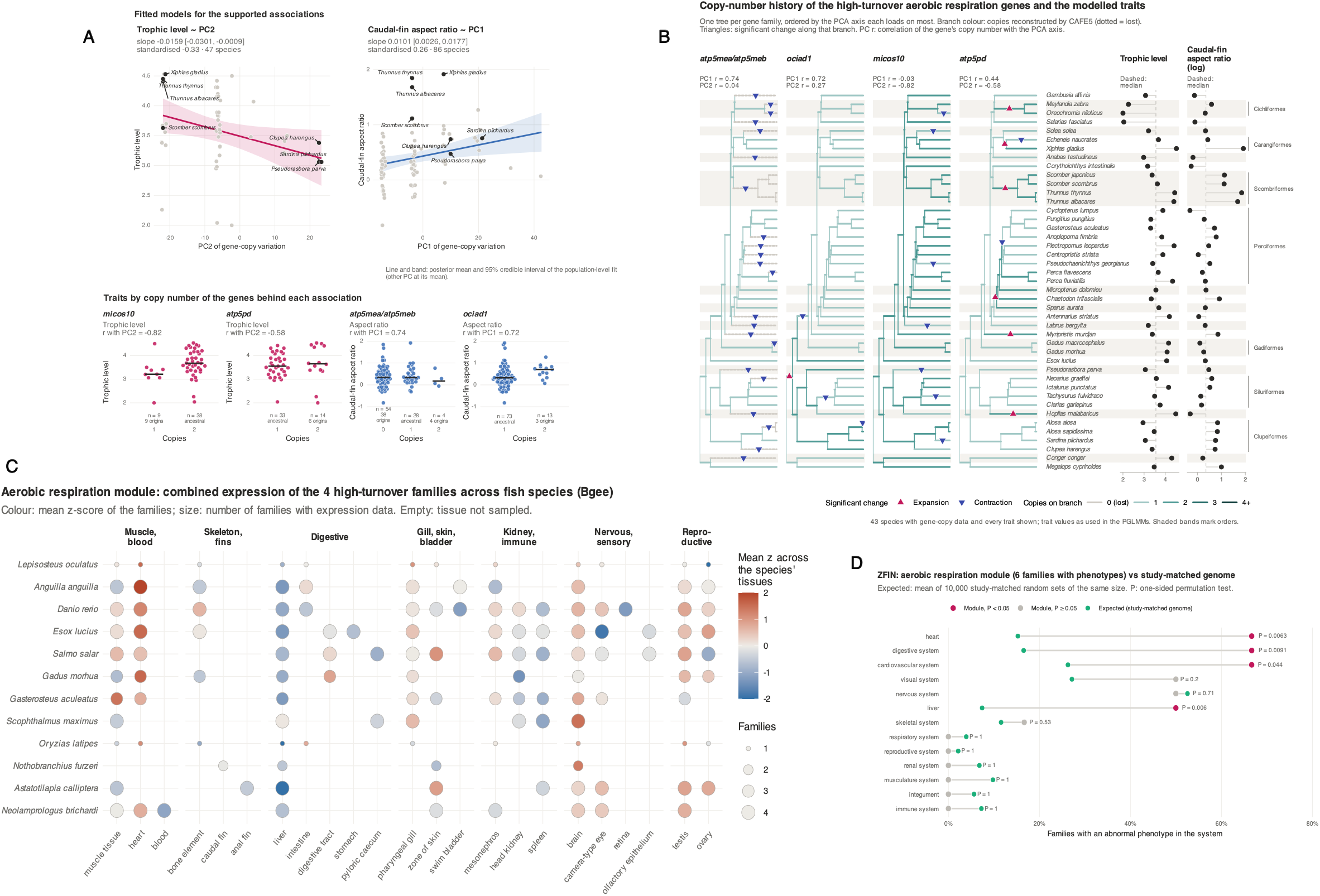
Variation in copy number of high turnover genes from the aerobic respiration module is associated with variation in trophic level and swimming performance. (A) Fitted relationship for the two supported associations whose 95% credible intervals excludes zero. Trophic level decreases along PC2 while Log caudal-fin aspect ratio increases along PC1. Lines and shaded bands show posterior mean and 95% credible interval of the population-level fit. The points are species. The slope and standardised values are obtained from the model output. The lower panel shows trait values grouped by copy number that correlate strongly (r >= 0.5) with the PC axes. Each point is a species and bars are median. For each group n = number of species, the number below denotes independent origin for that copy number, ancestral is the reconstructed state obtained from CAFE at the root of the species. For example, a single-copy configuration of micos10 arose in 9 lineages which have a lower median trophic level than species retaining the two copies. (B) The plot shows the history of copy-number gain and loss for species used in the PGLMM. The trees represent the phylogenetic tree and species used in the PGLMM. Copy numbers at the nodes were estimated from the ancestral state reconstruction in CAFE. Trait values are displayed as higher or lower relative to the median trait value across species. (C) Combined expression of the high-turnover genes across tissues and species. The z-scores were calculated across available tissue expression data for each species. (D) Comparing with a random set of genes that had a comparable number of studies reporting phenotypes (study-matched), we found that perturbation phenotypes of genes from the aerobic respiration module were more frequently found in the heart, digestive system, and liver. The dots show the relative proportions of each category. This plot shows these genes tend to affect the above systems at a higher rate than other genes from the genome.

The associated genes were expressed in tissues with substantial energetic demands. Across 11 teleost species and spotted gar, the genes showed high combined expression in the heart, pharyngeal gills, and brain (Fig 5C). These patterns are consistent with their known roles in mitochondrial organisation and oxidative metabolism (*50–53*). Moreover, data from ZFIN showed that perturbation of genes from the aerobic respiration module more frequently affected structures related to the heart and digestive system than any random set of genes (one-sided permutation test; 10,000 iterations; P < 0.05) (Fig 5D). Together, they show that these genes function in tissues that can have a plausible effect on the associated phenotypes.

## Discussion

Metamorphosis in teleosts is a transition involving the coordinated remodelling of multiple biological systems (*10*, *21*, *28*, *42*). The extent to which this transition is based on lineage-specific or on shared developmental frameworks has remained unclear. Here, we provide evidence that post-embryonic development in teleosts is governed by more than an assortment of independent, lineage-specific gene interactions. Instead, it is organised around a central genetic architecture that was already established early in teleost evolution.

In evo-devo there is an apparent tension between the idea of conservation and diversification. Can biological processes that stay conserved across species also contribute to the formation of variation? Understanding how a conserved system can incorporate evolutionary changes to generate phenotypic variation is therefore central to resolving this tension. Our results provide clues as to how this can be achieved. The conserved network is partitioned into two functional components: core cellular processes which share components with embryogenesis, and a remodelling partition that is unique to post-embryonic development. The former provides the molecular building blocks required for cells to proliferate, differentiate, and grow while the latter comprise programmes more directly involved in developmental remodelling and physiological maturation. This demonstrates how the same set of genes in an organism can be organised into different functional entities to carry out specific functions at specific times. This means that the underlying organisation of post-embryonic development is not between developmental and non-developmental genes, but between genetic organisation that enables developmental change and genetic programmes that execute tissue remodelling. This mechanism highlights the redeployment of conserved features.

The conserved network is particularly responsive to thyroid hormone (TH) induction. Response to thyroid hormone is a hallmark of metamorphic remodelling (*7*, *10*). Thus our result places the conserved network as an integral part of metamorphic remodelling and not merely sets of genes that change in expression during development. However, these modules did not respond uniformly to TH induction. The core partition was more responsive than the remodelling partition. In *Xenopus,* the initial response to TH induction is a general increase in cell proliferation and metabolism, followed by more tissue-specific changes tied to remodelling at later stages (*54*). A similar hierarchical response would explain why the broadly expressed core partition responds to a higher degree than the more tissue-specific remodelling partition. This difference in response suggests that there exists a form of regulatory fine tuning that can incorporate variation in the temporal-spatial activity of the individual modules without requiring the entire developmental programme to be replaced or rewired. This regulatory fine tuning can incorporate functional specification to the network allowing parts of it to respond selectively to ecological variation. In the manini ecological experiment, habitat explains a substantial portion of the variation in the remodelling partition, with most of it being focussed on the neural modules and peroxisome module. In contrast, the core cellular partition is less responsive to habitat variation and instead tracks developmental progression. This separation potentially highlights regulatory refinement, making it possible to restrict network response to only those components that would be beneficial under specific environmental conditions. Although more experimental evidence is required to validate this hypothesis, it is consistent with observations across teleosts of endocrine mediated responses to environmental stress (*55*, *56*), and environmentally induced phenotypic plasticity (*42*).

The conserved network contains genes with high rates of copy number turnover, indicating that its architecture is not evolutionarily static. Rather, the network has evolved through changes in gene copy number, a characteristic feature of biological network evolution (*57*). Such changes could modify gene dosage, allow paralogs to acquire specialised functions, or through gene loss, remove or simplify existing components (*58*, *59*). These mechanisms could modify a conserved developmental architecture while maintaining its overall integrity. The modules involved in neuronal functioning stand out in our results. They are specific to post-embryonic development, have conserved expression and tissue specificity across teleosts, yet have experienced high turnover in gene copy numbers and harbour extensive ecological variation. They contain genes which contribute to processes through which animals perceive and respond to their environment. For example, *grin2da/grin2db* encode glutamate receptor subunits whose diversification has contributed to establishing vertebrate synaptic plasticity and behaviour (*60*). The melanopsin genes *opn4a* and *opn4b*, which produce non-visual photopigments, have roles in maintaining circadian rhythms and modulating visual inputs (*61*). The modules also include *ahr1b,* which encodes a ligand-activated aryl hydrocarbon receptor. It is considered a vertebrate specific innovation and predicted to have an adaptive role in regulating xenobiotic metabolic enzymes (*62*). Studies across diverse groups of animals have shown that such ‘environment facing’ genes tend to also be enriched for adaptive molecular convergence (*63–65*), thereby making them key targets for adaptation in organisms-environments interactions. This suggests that despite retaining their conserved positions within the developmental network, variation genes in the neuronal modules can be potentially repurposed to enable functional innovation.

While the evolutionary genomic analysis provides a broad perspective on how the components underlying post embryonic development evolved, our phylogenetic models allow us to formulate specific hypotheses linking genomic variation to phenotypic diversity. Although these links do not by themselves distinguish causation from correlated evolutionary change, the clustering of distantly related species in similar regions of the phylogenomic space (PC1 and PC2 in Fig 5A) can indicate the existence of convergent evolutionary outcomes (*64*, *66–68*). This transforms the conserved network from a descriptive developmental pattern into a plausible substrate for phenotypic evolution. The result of the aerobic respiration module provides an effective example of this. The high-turnover genes in the aerobic respiration module functionally converge on mitochondrial structure and oxidative energy metabolism. *Micos10* encodes MIC10, a component of the MICOS complex that maintains crista junctions and inner mitochondrial membrane architecture, where oxidative phosphorylation occurs (*69*), (*50*, *51*). *Atp5meb* and *atp5pd* encode mitochondrial ATP synthase subunits and are directly linked to proton motive force-driven ATP production (*48*), (*52*). In contrast, *ociad1* regulates mitochondrial metabolism through electron transport chain Complex I and helps maintain the metabolic state required for progenitor-cell maintenance and differentiation ((*53*), (*70*)). These genes suggest that this module captures variation in mitochondrial organisation, aerobic ATP production, and cellular metabolic regulation. In teleosts, this module may be especially relevant because aerobic energy metabolism is tightly linked to swimming performance, and feeding strategy. Predatory species that rely on sustained swimming, such as tunas and swordfish, have high mitochondrial densities in red muscle, supporting prolonged activity and, in some cases, regional endothermy (*71–74*). Similarly, fishes specialised for sustained body-caudal-fin swimming often have elongate, streamlined bodies associated with higher oxygen demand (*75*). These links suggest that the aerobic respiration module may represent an important developmental and evolutionary axis connecting mitochondrial function, with diverse swimming modes and trophic strategies in teleosts.

## Conclusion

The post embryonic developmental network is conserved, yet evolvable. Most of the essential body plan is already established during embryogenesis, therefore variation in post-embryonic development might allow individual traits to vary without disrupting the overall organisation of the animal. This would make it better suited to accumulate adaptive variation. Furthermore, traits that would affect reproductive fitness of the adult are formed during this stage. While embryonic patterning may influence their establishment, studying it alone leaves a substantial portion of the developmental process, and how it can vary across ecologies unresolved. Our results therefore encourage greater focus towards the study of post embryonic remodelling across a wide group of organisms to better understand how conserved systems can be a seat of diversification.

## Materials and Methods

### Data collection

We used RNA-seq data from three previously published studies on clownfish (*28*), grouper (*21*), and manini post-embryonic development (*30*). For this study we generated data for zebrafish and goby. The zebrafish stages were based on descriptions in (*33*) and goby on (*31*). A detailed description of the methods regarding sample collection, RNA extraction, and sequencing can be found in the supplementary methods 1 in the supplementary material.

We used genomic data of 151 teleost fish species (including the above five), and spotted-gar *Lepiosteous oculatus* as an outgroup. Data were collected from the National Center for Biotechnology Information (NCBI) (*76*), Ensembl genome databases (*77*), and individual study sources. A list of genomes and their sources can be found in table S5. All genomes were highly contiguous, chromosomal level genomes with gene annotations available. For all analyses we used pruned versions of the rooted time-calibrated species tree from the Fish Tree of Life project (*36*).

### Orthology estimate

Orthologous relationships were inferred at the level of the last common ancestor (LCA) of the 152 fish species in the dataset using a rooted, time calibrated phylogenetic tree from the Fish Tree Of Life project (*36*). In this framework, each orthologous group (ortholog group) corresponds to genes derived from a single ancestral gene (or genes) present in the LCA of all the compared species. The annotated gene sets (proteomes) for the 152 species were preprocessed to filter the longest isoform per protein using the in-house python script. Orthology inference was performed with the standalone OrthoDB pipeline OrthoLoger v3.0.2 (*78*) using default settings. The OrthoLoger workflow first conducts all-against-all pairwise protein sequence comparisons between and within species, to detect best reciprocal hits across all the genes. These BRH relationships are then used as the basis for a graph-based clustering approach that begins with BRH triangulation and iteratively expands clusters to reconstruct ortholog groups, aiming to capture all genes that trace back to an ancestral gene(s) in the LCA. In zebrafish we identified 18,063 ortholog groups, 17,576 ortholog groups in clownfish, 18,662 ortholog groups in manini, 18,533 ortholog groups in grouper, and 18,378 ortholog groups in goby. To obtain the orthologous gene expression matrix we filtered out genes that were expressed across all stages in all the five species. This resulted in 12,700 ortholog groups accounting for ∼70% ortholog groups in each species. The excluded ortholog groups were involved in a limited number of processes related to cell signalling, immunity, and cell division, and functioning in a few cellular compartments (table S6).

### Network construction

To construct the network we first quantified RNA-seq libraries of the developmental time series using Kallisto v0.46.1 (*79*). We then normalised for library size using counts per million transformation (CPM) in edgeR (*80*) and filtered out genes that had >10 CPM expression in at least three replicates per stage. Next, we log-transformed CPM values with a pseudocount of 1e-5 to and used Co-expression Batch Reduction Adjustment (COBRA) to decompose the covariance structure of the time-series expression data. COBRA represents gene–gene covariance through an eigendecomposition, yielding a matrix of eigenvectors that describes the major axes of transcriptomic covariance across samples. Because our objective was to identify transcriptional programs spanning the complete developmental transition, this matrix was used as input for WGCNA (*81*) to construct developmental networks individually for each species. We used the *pickSoftThreshold* function to select the best power that attains the scale-free topology of biological networks (*81*). The power values were similar for all datasets (goby:12, the rest 13). We used a minimum module size of 50 and merged modules using the Dynamic Tree Cut approach with a threshold of 0.25. This resulted in modules with genes grouped together based on similar contributions across the major covariance axes of the developmental transcriptome. Therefore, the modules can be interpreted as covariance-defined transcriptional programs whose genes show coherent co-expression patterns across development (Fig 2A). To ensure that the modules we obtain represent biologically meaningful grouping of genes due to their activity during post-embryonic development, we constructed networks using permuted gene expression values and compared the network structures. The random networks failed to achieve scale free topology, have vastly different connectivities, and the modules do not show any preservation between species. Therefore, the modules from our original networks are a biologically meaningful representation of coordinated gene activity during post-embryonic development, and not simply transcriptomic grouping of housekeeping genes. Full details in supplementary methods 2.

### Obtaining the conserved co-expression network

We estimated module preservation between module pairs which measures how well network properties are conserved between a reference and a test network (*37*). The *Z_summary_* score is used to denote the degree of preservation with scores above 10 typically used as evidence of high preservation and scores below 2 as evidence of no preservation (*37*). To get more reliable estimates we use more stringent thresholds to categorise preservation, with *Z_summary_ >* 12 as evidence of high preservation, *Z_summary_ <* 12 as evidence for low to medium preservation, and *Z_summary_ <* 6 as no evidence for preservation. To estimate module memberships we used the Jaccard similarity index. For each pair of species, we compared every module in species A with every module in species B. For each module pair (Mi,Nj) we calculated the Jaccard similarity of gene membership, J=Mi Nj Mi Nj, and estimated the overlap between member genes using a one-sided Fisher’s exact test. Only the module pairs with evidence for significantly higher overlap (Benjamini-Hochberg corrected P < 0.05) were retained. Among these significant pairs we then ranked them by Jaccard similarity and kept the top 30%.

Using higher thresholds (top 25% etc) reduced the number of overlapped genes below 10% of the genes used to construct the network, whereas, using lower thresholds introduced noise and made the modules hard to interpret. Therefore, we select the 30% threshold as it ensures we retain only the modules that have the highest proportion of co-expressed genes between each species pair, reducing noise and false positives. After repeating this procedure for all species pairs, we intersected the retained gene sets across all comparisons to obtain a cross-species set of ‘conserved network’ genes. These genes are consistently co-expressed within significantly overlapping modules in all five species during the post-embryonic transition. GO term enrichment was carried out using a hypergeometric test implemented via the clusterProfiler package (https://github.com/YuLab-SMU/clusterProfiler).

### Comparing embryonic and post-embryonic zebrafish networks

We used an embryonic developmental time series from (*38*) and restricted the data to 15 embryonic stages (1 cell to Long-pec). We used the exact same zebrafish genes that were used to construct the original zebrafish post-embryonic developmental network. Of the 12,700 genes used to construct the original post-embryonic network, 8395 passed the expression filter in the embryonic time series. Therefore, we re-constructed the networks using these 8395 genes. The new networks were built using the exact same WGCNA setting with soft-threshold powers between 8-10 tested for sensitivity. The module preservation was measured in both directions by running for 200 iterations. Module preservation was estimated pairwise for all modules in both directions. For the six network comparison (zebrafish embryonic network x 5 species post-embryonic network), genes overlapping between the networks were used.

### Thyroid hormone induction and in-situ experiment

The data for the T3 and MPI treatments was obtained from a previous study (*28*). A brief description of the experiment is provided in supplementary methods 3. We used edgeR (*80*) quasi-likelihood tests (FDR < 0.05) to estimate differential expression (DE). Using two backgrounds we tested for enrichment of DE genes in the conserved network. The first background was all the 12,700 ortholog groups used to construct the clownfish network but not in the conserved network. The second was the entire transcriptome. We also split the background and conserved network into expression matched bins and compared the rates of DE genes in the background and conserved network. This approach allows us to account for the high genomic conservation in the orthologous groups and the higher expression levels of genes from the conserved network.

### The data for the manini in-situ experiments were obtained from (*30*). Settlement-stage

*A. triostegus* larvae, which were fully transparent and classified as stage S0, were collected along the northeast coast of Moorea Island in French Polynesia. Following capture, larvae were assigned to two experimental groups. One group was transferred to aquaria at the CRIOBE Marine Research Station (this was used for the developmental network), whereas the second group was placed in in-situ cages distributed across the three nursery habitats. Fish were sampled 12 hours (S1), 3 days (S2), and 8 days (S4) after capture. The variation in environmental features and metabolic features of the fish across habitats was verified in the original manuscript (*30*). For each gene, we modelled expression as a function of developmental stage (gene expression ∼ stage) or habitat and developmental stage (gene expression ∼ stage + habitat). We then measured the gain in proportion of variance explained by habitat as a reduction in residual sum of squares when adding the habitat term, divided by the total sum of squares. This model allowed us to obtain expression variance due to a developmental stage component (corresponding to developmental progression shared across habitats), and a habitat component. The median variances of gene sets were compared against an expression-matched background of orthologous genes not in the conserved network. with their median variance explained centered around 1. To account for unbalanced sampling (beach rock habitat had only 3 replicates of S3, and no S4) we reran the analysis using several designs and observed consistent results for each design. Habitat explained less variance in the conserved network than the background, and habitat consistently explained more variance in the remodelling partition versus the core (table S7). Using a leave-one-out approach we recomputed the difference between the partition variances to determine whether a few modules were driving the high habitat variance of the remodelling partition. The results confirmed that, rather than a single module, the neural modules together were driving the difference in habitat variance between the two partitions (table S8).

### Evolutionary analysis

Out of the 152 species for which we generated ortholog groups, we excluded teleost lineages with additional lineage-specific WGDs because their genomes introduce extra paralogy, lineage-specific duplicate retention and loss, and extended rediploidisation dynamics, which can bias comparative estimates of copy-number evolution and regulatory divergence (*82*, *83*). Our evolutionary analyses therefore focused on genomes sharing the same duplication background. The final list included 134 species and can be found in table S9. To represent the goby, we used the single branch leading to *Rhinogobius brunneus* as its phylogenetic proxy. We processed sequences per ortholog group prior to running evolutionary codon models. We used the CDSKIT v0.91 tool (https://github.com/kfuku52/cdskit) to make the sequences in-frame. Next, using TRANSeq v6.6.0 (*84*) we translated the coding sequences into protein sequences followed by alignment using MAFFT v7.481 with the ‘auto’ option (*85*). We then performed a translation alignment using TRANALIGN v6.6.0 (*84*) and trimmed poorly aligned sequences using TRIMAL v1.4.1(*86*) with the ‘*ignorestopcodon*’ and ‘*automated1*’ options. Finally, we masked and removed ambiguous sites in the alignment using the ‘*mask*’ and ‘*hammer*’ functions of CDSKIT v0.91.

We used IQ-TREE v2.0.3 (*87*) with the MPF model, and 1000 ultra-fast optimised bootstrapping replicates (‘*bnni*’ option), to construct gene trees. We rooted and reconciled the gene trees with the species trees using a combination of GeneRax v1.1.0 (*88*) and NWKIT v0.10.0 (https://github.com/kfuku52/nwkit).

We used several molecular evolution models to test for signatures of selection at the coding-sequence level. We first used a standard MG94 selection model (*89*) to get a global estimate of non-synonymous to synonymous substitutions (dN/dS) across the entire phylogenetic tree. Next, we estimated dN/dS on a per-site basis using the FUBAR (Fast Unconstrained Bayesian Approximation) approach (*90*), which classifies individual sites as evolving under purifying selection, diversifying selection, or as having insufficient statistical support for assignment to either regime. Lastly, we ran the BUSTED (Branch-Site Unrestricted Statistical Test for Episodic Diversification) model to test for selection at the gene level (*91*). BUSTED fits a codon model with multiple rates classes to identify selection acting on at least one codon site along at least one branch of the phylogeny. These models were implemented using the HyPhy package (*92*).

We used CAFE5 v5.0 (*44*) to estimate evolutionary dynamics in copy-number evolution. First we ran a single-rate error corrected base model and a multi-rate model. The error model accounts for any non-biological factors such as genome sequencing and coverage differences, as well as gene family count estimation errors (*44*). We compared model fits using negative log-likelihood scores. Twenty-four of the 25 conserved modules were included in the analysis. The sterol synthesis module was excluded because its gene-count matrix was too sparse for reliable analysis.

### Phylogenetic generalised linear mixed model (PGLMM)

We ran PGLMMs using the brms v2.23.0 R package (*93*). Phenotypic trait data was obtained from the FishLife project (*46*). The FishLife project is a collaborative international project that collects life-history data followed by expert curation allowing seamless integration into comparative methods workflows (*46*). The data was accessed through the FishLife v3.1.0 R package (*46*). Transformed traits were used as the response variable. We use the gene family count matrix of the high turnover gene families to compute a phylogenetic PCA (pPCA) using phytools v2.5.2 (*94*). The count matrix was scaled prior to computing the pPCA. We then use the scores of the first two components of the pPCA as predictor variables for the PGLMM. This allows us to effectively capture the multi dimensional variation caused by changes in multiple gene copies without over parameterising the model, thereby preventing the classic *n*x*p* problem common in comparative genomics (*95*). It is important to note that the pPCA scores are still in the original phylogenetic species space, and the non-independence of species still has to be accounted for (*47*). Therefore, we model the phylogenetic effect as a group level (random) effect using the covariance structure from the phylogenetic tree. Only species that were present in the phylogenetic tree were used. A skew normal distribution family was used for continuous data while a categorical distribution family was used for categorical traits. All models were run for 4000 iterations with 1000 warmups. Default priors were used in all models. For the body shape model we used a slightly tighter prior for the phylogenetic term. This produced better convergence as it prevented the markov chain from deviating into remote regions of the posterior space. All model convergence was determined using the multiple diagnostics (Rhat=1, Bulk_ESS > 2000, posterior distributions checks).

### Gene expression analysis from Bgee and perturbation phenotypes from Zebrahub

Multi-tissue and multi species gene expression analyses were carried out using data from the Bgee database (*49*). We compile data for 13 fish species (*Anguilla anguilla*, *Astyanax mexicanus, Astatotilapia calliptera, Danio rerio, Esox lucius, Gasterosteus aculeatus, Gadus morhua, Lepiosteous oculatus, Neolamprologus brichardi, Nothobranchius furzeri, Oryzias latipes, Salmo salar,* and *Scophthalmus maximus*). We only selected genes that were reported “present” and compared gene expression in a maximum of 23 tissues in only the adult stage. Using the zebrafish genes as the query, we obtained orthologs for the other species using Bgee’s API calls. We used log2(TMP+1) gene expression estimates, and for each ortholog group the copies were summed in each species. A z-score was calculated across a species’ sampled tissue and averaged over a module’s families per tissue.

We use the tissue-specificity index tau (*96*) to estimate tissue. For each species, tau was calculated for every gene (not ortholog group) on log2(TPM+1) across tissues sampled in that species. For each module, we used the median tau of all the genes in the module as the representative value. This was compared against the median tau of an expression-matched equal-sized null set of random genes in that species. The final statistic was △tau = module median - median of the random null, where below 0 means a module is more orally expressed against the null and above 0 means it is more specific than the null. We computed Benjamini-Hochberg adjusted empirical P-values for each module-null comparison. *Nothobranchius furzeri* was not used to compute tau due to its low tissue distribution.

For each module we searched for perturbation data for module genes in the ZFIN database and calculated the number of unique phenotypic terms and the biological system affected with the perturbation. We then compared the distribution of affected systems with a distribution of equal-studied random genes. A gene that has hundreds of studies associated with it will by design have more reported phenotypes. Therefore the equal-study null ensures we compared genes with similar research depth. We then used a one-sided permutation test to evaluate whether perturbation of module genes were enriched in specific tissues systems. The tissue system annotations were based on the zebrafish anatomy ontology.

## Author contribution

AB conceived the project, obtained the funding, carried out all the analyses, and wrote the manuscript. GC performed the orthology estimate. SM carried out all the molecular biology experiments. MH, KM, YY, YG, and VL provided samples. VL and MRR provided in-depth feedback on the manuscript. All authors approved the manuscript.

## Data availability

Sequence data for zebrafish have been deposited in NCBI under the BioProject PRJNA1456083, data for goby is deposited under PRJNA1457023. All code, data, analyses, and outputs have been deposited in Zenodo. 10.5281/zenodo.23730920.

