## Supplementary material for "A post-embryonic specific and ecologically selective network underlies metamorphic remodelling in teleosts": Fig S

Agneesh Barua[1,2,*] Giulia Campli[1,2] Marcela Herrera[3], Saori Miura[3], Yo Y. Yamasaki[4], Ken Maeda[3], Yann Gibert[5], Vincent Laudet[3,6] Marc Robinson-Rechavi[1,2]

[1]Department of Ecology and Evolution, University of Lausanne, Lausanne, Switzerland.
[2]Evolutionary Bioinformatics, Swiss Institute of Bioinformatics, Lausanne, Switzerland.
[3]Marine Eco-Evo-Devo Unit, Okinawa Institute of Science and Technology Graduate University, Onna son, Okinawa, Japan.
[4]Ecological Genetics Laboratory, Department of Genomics and Evolutionary Biology, National Institute of Genetics, Yata 1111, Mishima, Shizuoka 411-8540, Japan
[5]Indiana University, School of Medicine.
[6]Marine Research Station, Institute of Cellular and Organismic Biology (ICOB), Academia Sinica, 23-10, Dah-Uen Rd., Jiau Shi, I-Lan 262, Taiwan

*Corresponding author


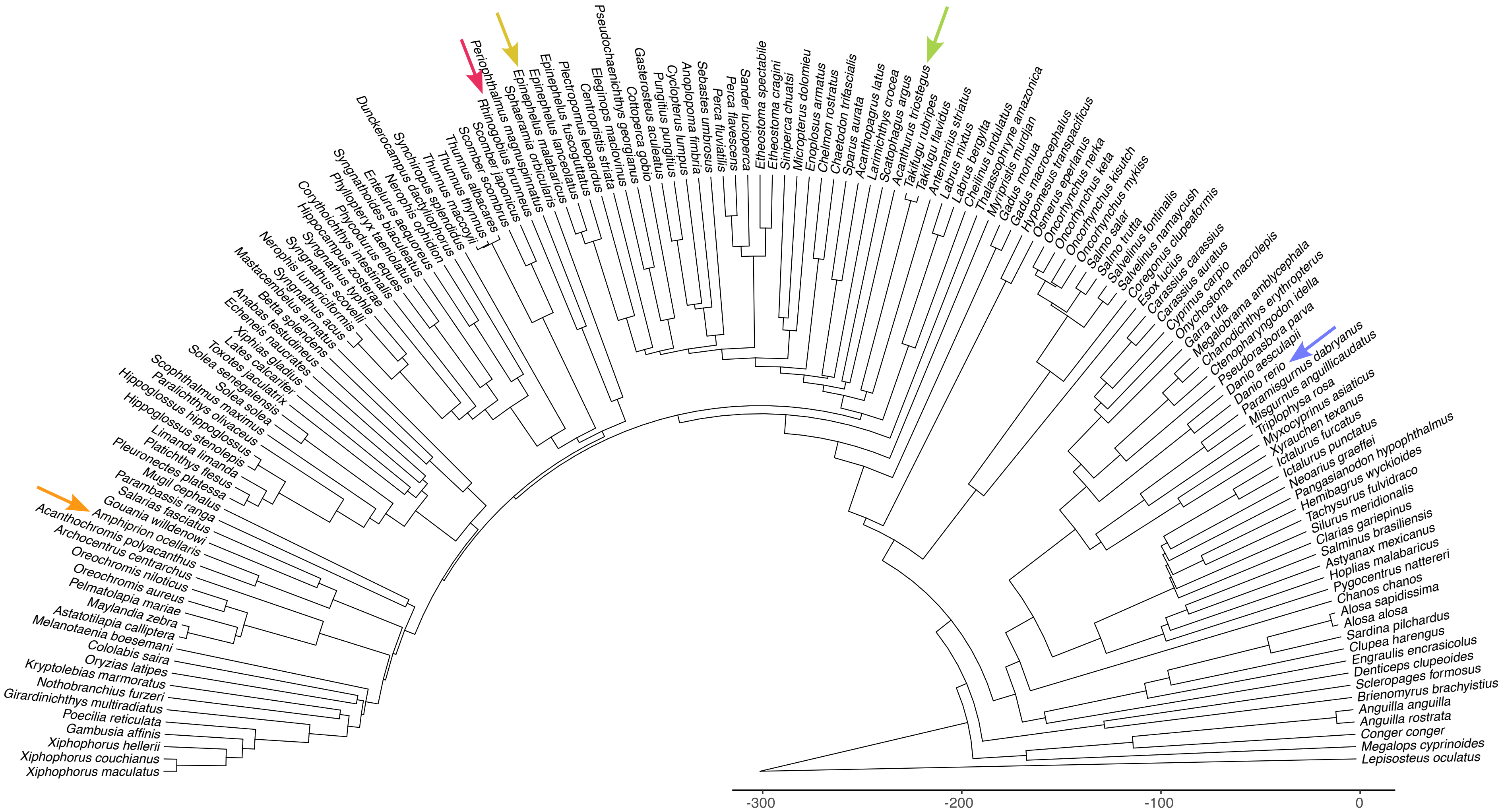

Xiphophorus maculatus
Xiphophorus couchianus
Xiphophorus hellerii
Gambusia affinis
Poecilia reticulata
Girardinichthys multiradiatus
Nothobranchius furzeri
Kryptolebias marmoratus
Oryzias latipes
Cololabis saira
Melanotaenia boesemani
Astatotilapia calliptera
Maylandia zebra
Pelmatolapia mariae
Oreochromis aureus
Oreochromis niloticus
Archocentrus centrarchus
Acanthochromis polyacanthus
Amphiprion ocellaris
Gouania willdenowi
Salarias fasciatus
Parambassis ranga
Mugil cephalus
Pleuronectes platessa
Platichthys flesus
Limanda limanda
Hippoglossus stenolepis
Hippoglossus hippoglossus
Paralichthys olivaceus
Scophthalmus maximus
Solea solea
Solea senegalensis
Toxotes jaculatrix
Lates calcarifer
Xiphias gladius
Echeneis naucrates
Anabas testudineus
Betta splendens
Mastacembelus armatus
Syngnathus acus
Nerophis lumbriciformis
Syngnathus scovelli
Syngnathus typhle
Hippocampus zosterae
Corythoichthys intestinalis
Phycodurus eques
Phyllopteryx taeniolatus
Syngnathoides biaculeatus
Entelurus aequoreus
Nerophis ophidion
Dunckerocampus dactyliophorus
Synchiropus splendidus
Thunnus maccoyii
Thunnus thynnus
Thunnus albacares
Scomber scombrus
Scomber japonicus
Rhinogobius brunneus
Periophthalmus magnuspinnatus
Sphaeramia orbicularis
Epinephelus malabaricus
Epinephelus lanceolatus
Epinephelus fuscoguttatus
Plectropomus leopardus
Centropristis striata
Eleginops maclovinus
Pseudochaenichthys georgianus
Cottoperca gobio
Gasterosteus aculeatus
Pungitius pungitius
Cyclopterus lumpus
Anoplopoma fimbria
Sebastes umbrosus
Perca fluviatilis
Perca flavescens
Sander lucioperca
Etheostoma spectabile
Etheostoma cragini
Siniperca chuatsi
Micropterus dolomieu
Enoplosus armatus
Chelmon rostratus
Chaetodon trifascialis
Sparus aurata
Acanthopagrus latus
Larimichthys crocea
Scatophagus argus
Acanthurus triostegus
Takifugu rubripes
Takifugu flavidus
Antennarius striatus
Labrus mixtus
Labrus bergylta
Cheilinus undulatus
Thalassophryne amazonica
Myripristis murdjan
Gadus morhua
Gadus macrocephalus
Hypomesus transpacificus
Osmerus eperlanus
Oncorhynchus nerka
Oncorhynchus keta
Oncorhynchus kisutch
Oncorhynchus mykiss
Salmo salar
Salmo trutta
Salvelinus fontinalis
Salvelinus namaycush
Coregonus clupeaformis
Esox lucius
Carassius carassius
Carassius auratus
Cyprinus carpio
Onychostoma macrolepis
Garra rufa
Megalobrama amblycephala
Chanodichthys erythropterus
Ctenopharyngodon idella
Pseudorasbora parva
Danio aesculapii
Danio rerio
Paramisgurnus dabryanus
Misgurnus anguillicaudatus
Triplophysa rosa
Myxocyprinus asiaticus
Xyrauchen texanus
Ictalurus furcatus
Ictalurus punctatus
Neoarius graeffei
Pangasianodon hypophthalmus
Hemibagrus wyckioides
Tachysurus fulvidraco
Silurus meridionalis
Clarias gariepinus
Salminus brasiliensis
Astyanax mexicanus
Hoplias malabaricus
Pygocentrus nattereri
Chanos chanos
Alosa sapidissima
Alosa alosa
Sardina pilchardus
Clupea harengus
Engraulis encrasicolus
Denticeps clupeoides
Scleropages formosus
Brienomyrus brachyistius
Anguilla anguilla
Anguilla rostrata
Conger conger
Megalops cyprinoides
Lepisosteus oculatus
-300
-200
-100
0

**Fig S1: Rooted time-callibrated teleost speices tree.** Phylogenetic tree of 151 teleost fish species and out group spotted-gar. The x-axis shows time in million years ago, with the root age ~300 million years. Orthologous relationships were inferred at the level of the last common ancestor (LCA) the 152 species. The arrows show the five species for which developmental time-series data was collected. The wide phylogenetic breadth ensure we capture genetic interactions that have been conserved since their LCA around 200 million years ago. This is pruned version of the tree form The Fish Tree of Life project. (Rabosky *et al*. 2018).

### Example 1

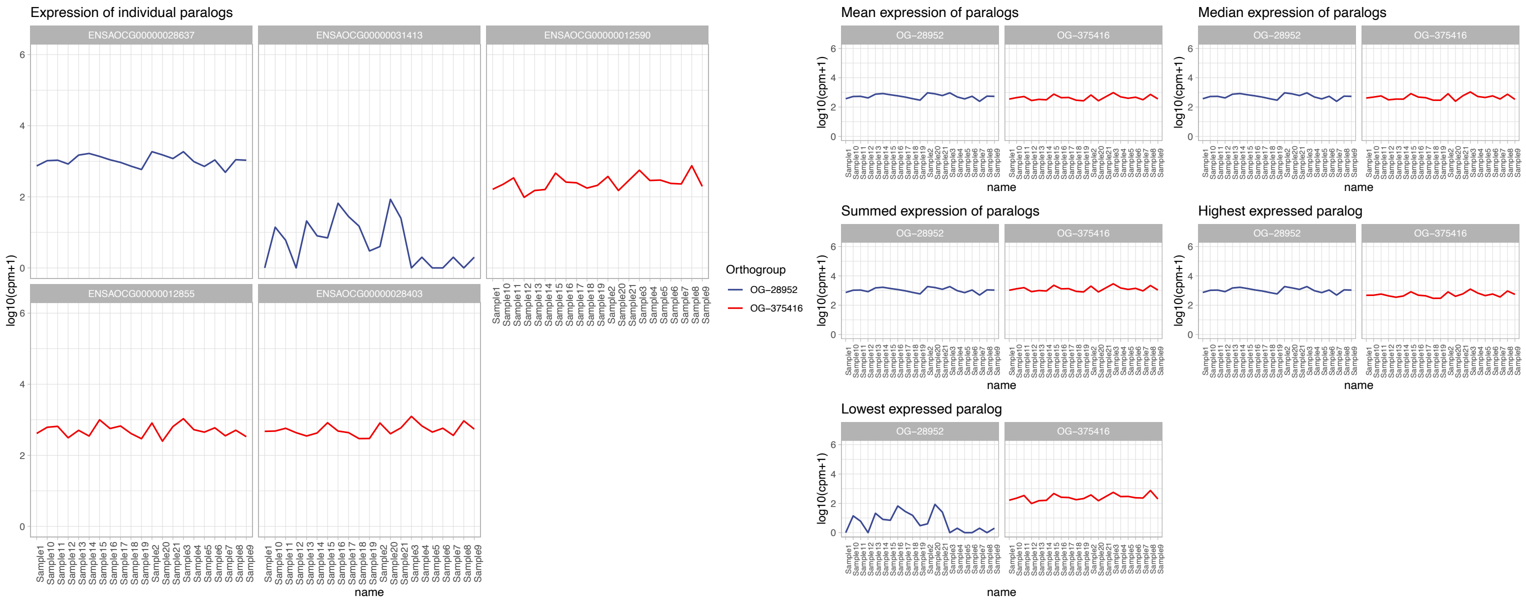


### Example 2

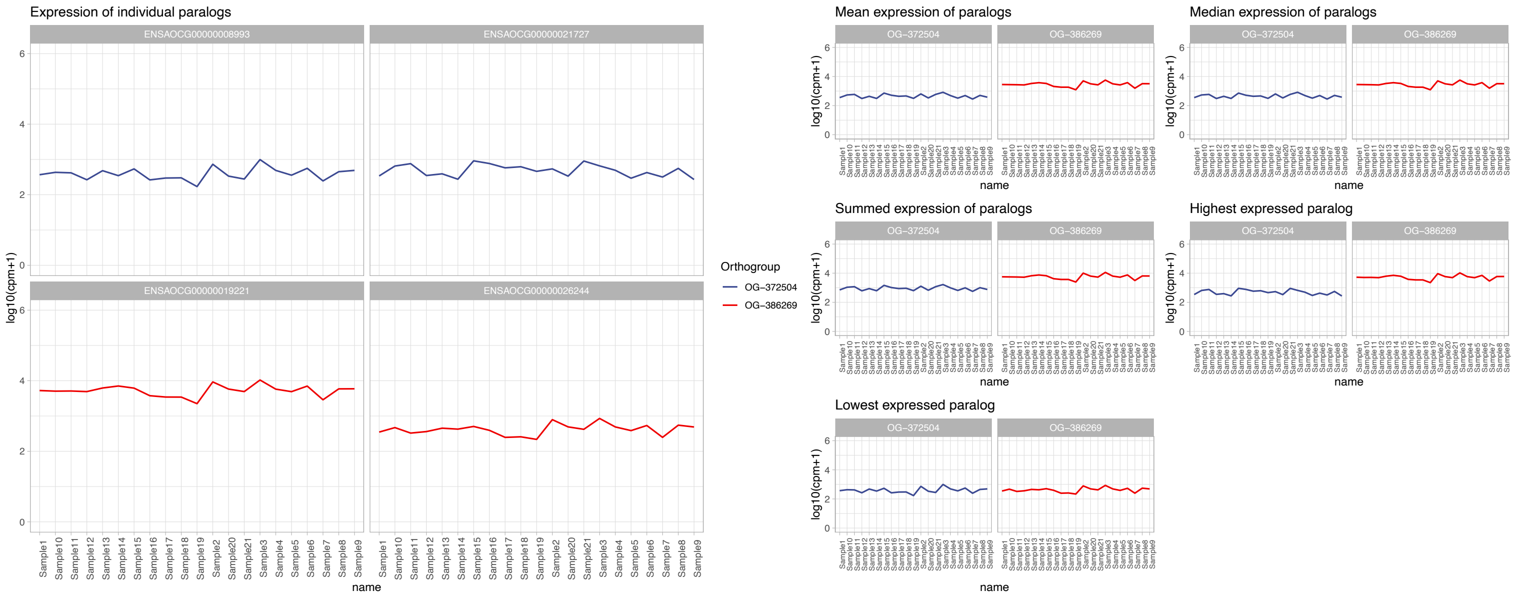


### Example 3

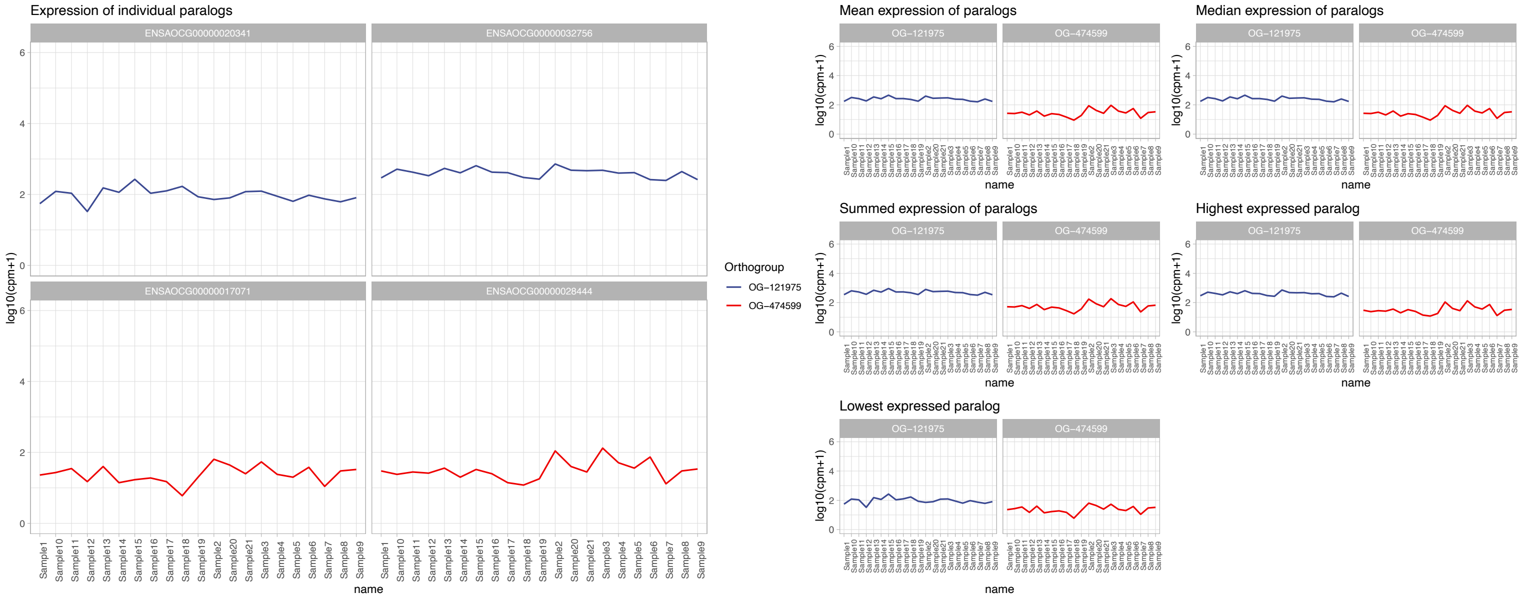

**Fig S2: Comparisons of summary statistics to quantify paralog expression.** All three examples show gene copies in different ortholog groups from clownfish. In example 1, there are two paralogous copies in the blue ortholog group, and three copies in the red ortholog group. The plots show how different summary statistics like mean, median, sum, and min/max expression for each copy all capture similar expression dynamics across development. Example 1 shows how one of the blue copies has a different expression profile as compared to the other copy from the same ortholog group. However, its average expression across development is magnitudes lower than the other, highly expressed copy. Furthermore, the highly expressed copy typical predominates the trend in expression which is similar to the other summaries of mean, median, and summed expression. Therefore, in our study we use the highest expressed copy as representative of expression level per ortholog group.

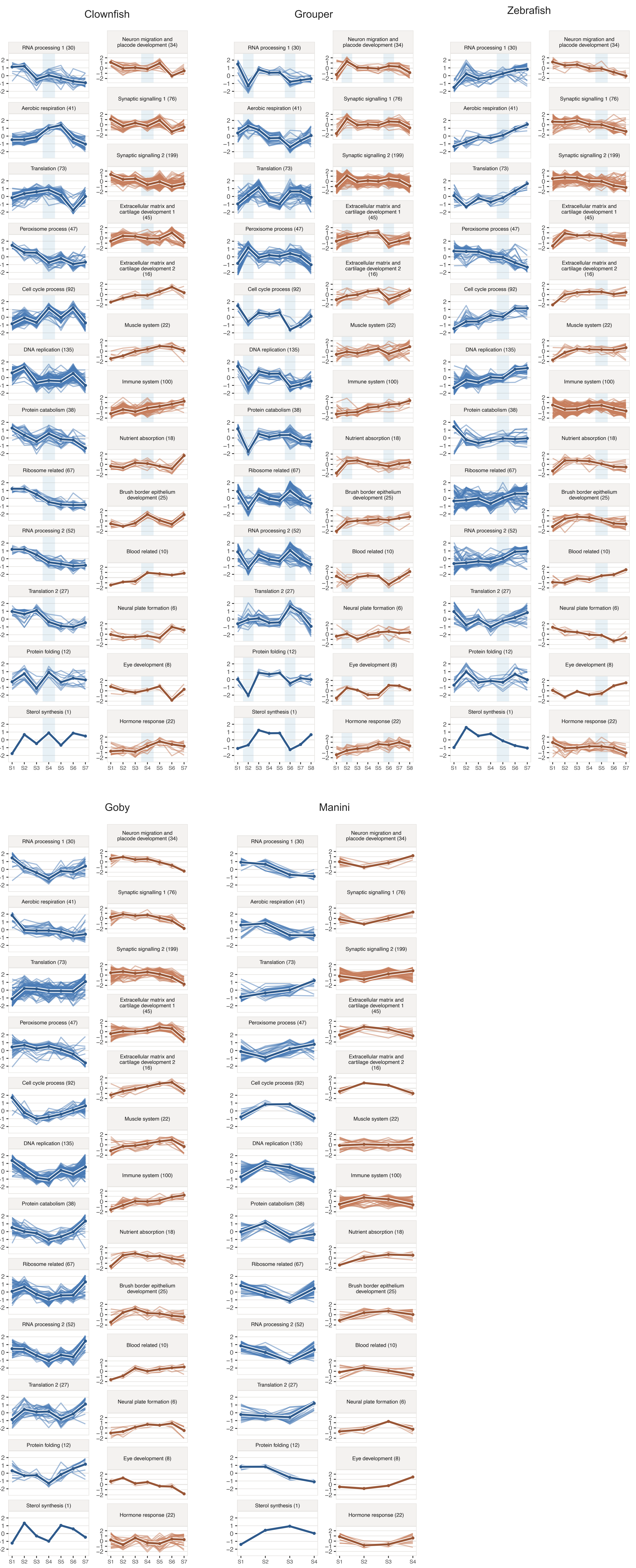
Clownfish
Grouper
Zebrafish
Goby
Manini
RNA processing 1 (30)
Aerobic respiration (41)
Translation (73)
Peroxisome process (47)
Cell cycle process (92)
DNA replication (135)
Protein catabolism (38)
Ribosome related (67)
RNA processing 2 (52)
Translation 2 (27)
Protein folding (12)
Sterol synthesis (1)
Neuron migration and placode development (34)
Synaptic signalling 1 (76)
Synaptic signalling 2 (199)
Extracellular matrix and cartilage development 1 (45)
Extracellular matrix and cartilage development 2 (16)
Muscle system (22)
Immune system (100)
Nutrient absorption (18)
Brush border epithelium development (25)
Blood related (10)
Neural plate formation (6)
Eye development (8)
Hormone response (22)
2 1 0 −1 −2
S1 S2 S3 S4 S5 S6 S7 S8

**Fig S3: Expression trajectories of the homologous modules of the conserved network.** Results for the expression of 1196 ortholog groups making up the 25 conserved module sets. Each panel comprises genes from different ortholog groups that were placed into a homologous module. The blue vertical bars denote peaks of thyroid hormone levels. The annotations of the homologous modules are obtained from GO enrichment.

#### Adult-tissue expression of the 25 conserved modules across fish species (Bgee)

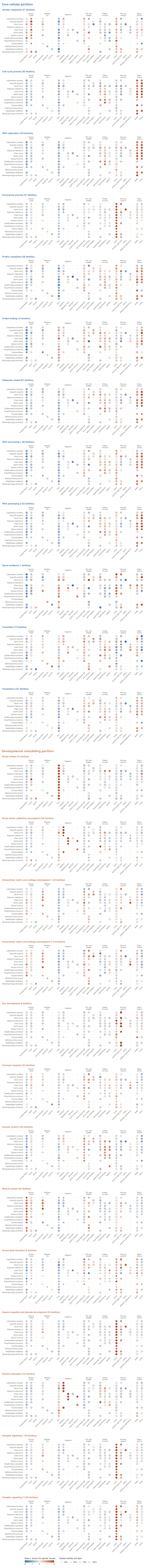


Colour: mean z-score of the module's families in each tissue: z of log2(TPM + 1) across the adult tissues sampled in each species, copies of a family summed.
Size: share of the module's families with expression data in that species. Empty: tissue not sampled, or no data.
Species in phylogenetic order. Outside zebrafish, genes are placed with Bgee orthologues of the zebrafish genes.

**Fig S4: Expression patterns of the homologous modules across teleosts.** Plots showing the combined expression of the modules across tissues in several species of teleosts and gar. The expression levels are plotted as z-scores computed across available tissues in each species. The numbers in brackets are the number of ortholog groups in each module. The size of the circles denote the percentage of the module with data.

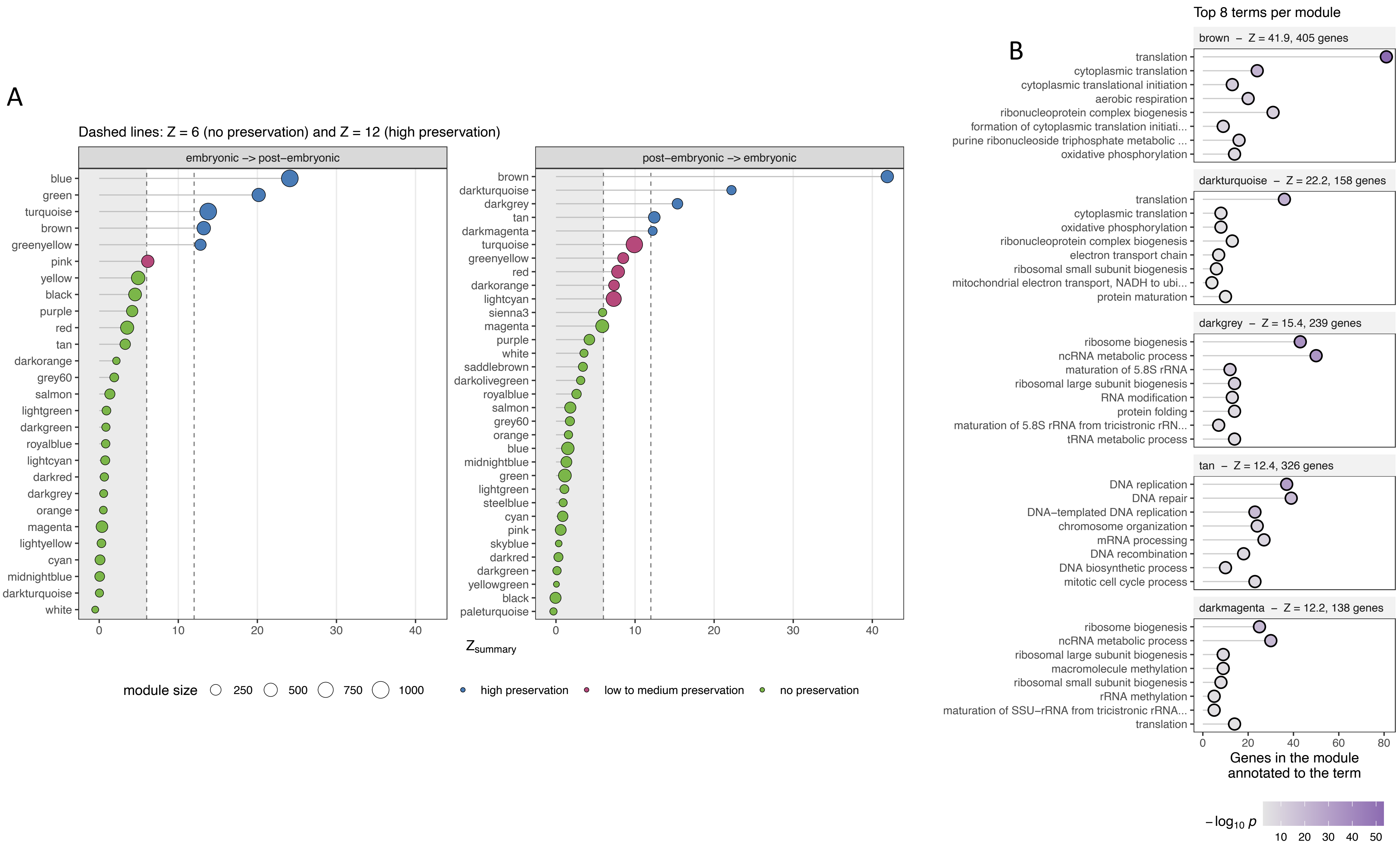

A
Dashed lines: Z = 6 (no preservation) and Z = 12 (high preservation)
embryonic -> post-embryonic
blue
green
turquoise
brown
greenyellow
pink
yellow
black
purple
red
tan
darkorange
grey60
salmon
lightgreen
darkgreen
royalblue
lightcyan
darkred
darkgrey
orange
magenta
lightyellow
cyan
midnightblue
darkturquoise
white
post-embryonic -> embryonic
brown
darkturquoise
darkgrey
tan
darkmagenta
turquoise
greenyellow
red
darkorange
lightcyan
sienna3
magenta
purple
white
saddlebrown
darkolivegreen
royalblue
salmon
grey60
orange
blue
midnightblue
green
lightgreen
steelblue
cyan
pink
skyblue
darkred
darkgreen
yellowgreen
black
paleturquoise
0
10
20
30
40
$Z_{summary}$
module size
250
500
750
1000
high preservation
low to medium preservation
no preservation
B
Top 8 terms per module
brown – Z = 41.9, 405 genes
translation
cytoplasmic translation
cytoplasmic translational initiation
aerobic respiration
ribonucleoprotein complex biogenesis
formation of cytoplasmic translation initiati...
purine ribonucleoside triphosphate metabolic ...
oxidative phosphorylation
darkturquoise – Z = 22.2, 158 genes
translation
cytoplasmic translation
oxidative phosphorylation
ribonucleoprotein complex biogenesis
electron transport chain
ribosomal small subunit biogenesis
mitochondrial electron transport, NADH to ubi...
protein maturation
darkgrey – Z = 15.4, 239 genes
ribosome biogenesis
ncRNA metabolic process
maturation of 5.8S rRNA
ribosomal large subunit biogenesis
RNA modification
protein folding
maturation of 5.8S rRNA from tricistronic rRN...
tRNA metabolic process
tan – Z = 12.4, 326 genes
DNA replication
DNA repair
DNA-templated DNA replication
chromosome organization
mRNA processing
DNA recombination
DNA biosynthetic process
mitotic cell cycle process
darkmagenta – Z = 12.2, 138 genes
ribosome biogenesis
ncRNA metabolic process
ribosomal large subunit biogenesis
macromolecule methylation
ribosomal small subunit biogenesis
rRNA methylation
maturation of SSU-rRNA from tricistronic rRNA...
translation
0
20
40
60
80
Genes in the module annotated to the term
$-\log_{10} p$
10
20
30
40
50

**Fig S5: Preservation statistics of modules between post-embryonic developmental network and embryogenesis network in zebrafish**. (A) For both comparison majority of the modules are not preserved. The five modules that are preserved in each comparison belong to the core cellular partition. (B) Top 8 GO terms enriched in each of the five post-embryonic network module that are preserved in embryogenesis (these are the same across both comparisons).

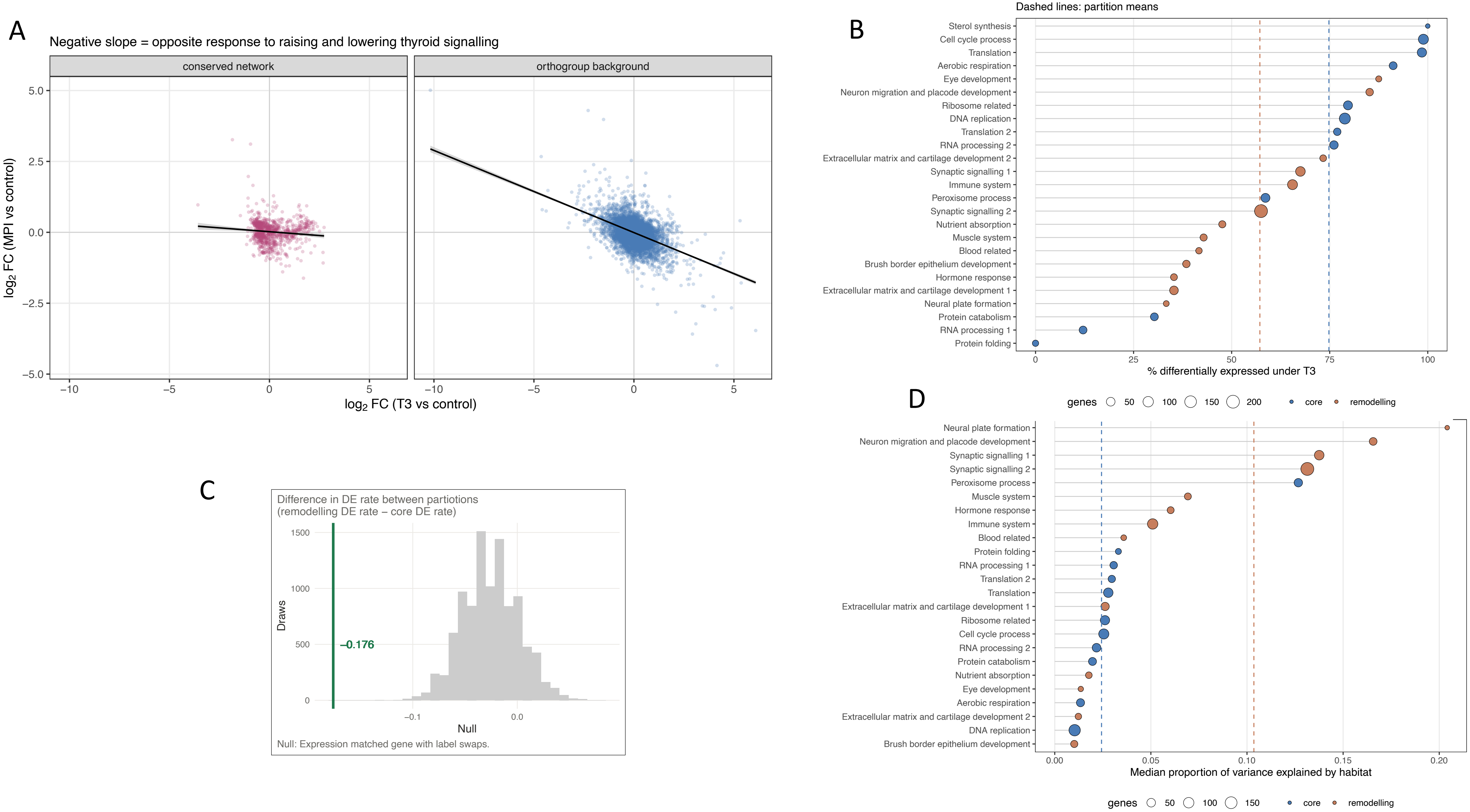
A
Negative slope = opposite response to raising and lowering thyroid signalling
conserved network
orthogroup background
log2 FC (MPI vs control)
log2 FC (T3 vs control)
5.0
2.5
0.0
−2.5
−5.0
−10
−5
0
5
B
Dashed lines: partition means
Sterol synthesis
Cell cycle process
Translation
Aerobic respiration
Eye development
Neuron migration and placode development
Ribosome related
DNA replication
Translation 2
RNA processing 2
Extracellular matrix and cartilage development 2
Synaptic signalling 1
Immune system
Peroxisome process
Synaptic signalling 2
Nutrient absorption
Muscle system
Blood related
Brush border epithelium development
Hormone response
Extracellular matrix and cartilage development 1
Neural plate formation
Protein catabolism
RNA processing 1
Protein folding
0
25
50
75
100
% differentially expressed under T3
genes 50 100 150 200
core
remodelling
C
Difference in DE rate between partiotions
(remodelling DE rate − core DE rate)
Draws
1500
1000
500
0
−0.176
−0.1
0.0
Null
Null: Expression matched gene with label swaps.
D
Neural plate formation
Neuron migration and placode development
Synaptic signalling 1
Synaptic signalling 2
Peroxisome process
Muscle system
Hormone response
Immune system
Blood related
Protein folding
RNA processing 1
Translation 2
Translation
Extracellular matrix and cartilage development 1
Ribosome related
Cell cycle process
RNA processing 2
Protein catabolism
Nutrient absorption
Eye development
Aerobic respiration
Extracellular matrix and cartilage development 2
DNA replication
Brush border epithelium development
0.00
0.05
0.10
0.15
0.20
Median proportion of variance explained by habitat
genes 50 100 150
core
remodelling

**Fig S6: Contrasting expression patterns between T3 and MPI (Goitrogen) treatments and statistics across the two conserved network partitions**. (A) The plot shows the relative difference in fold change of the same genes between the two treatments. Compared tot eh orthogroup background (Spearman rho = -0.33, -0.35 respectively; P= $2.2x10^{-16}$), the conserved network shows less contrasting expression in the two treatments (Spearman rho = -0.14; P= $2.2x10^{-16}$) demonstrating that the conserved network responds asymetrically to exogenous thyroid hormone treatment. (B) Genes of the core partitons were more frequently differentially expressed than the remodelling partition. (C) Differenes in rates of diferentially expressed genes belonging to the two partitions in the T3 treatment. Median $Core_{log2FC}$ = 0.539, median $Remodelling_{log2FC}$ = 0.387, differnce = -0.178. Difference is compared against expression matched null sets with labels shuffled. (D) The modules with the highest variance explained by habitat are neural modules belonging to the remodelling partition, and module involved in peroxisome process belonging to the core partition.

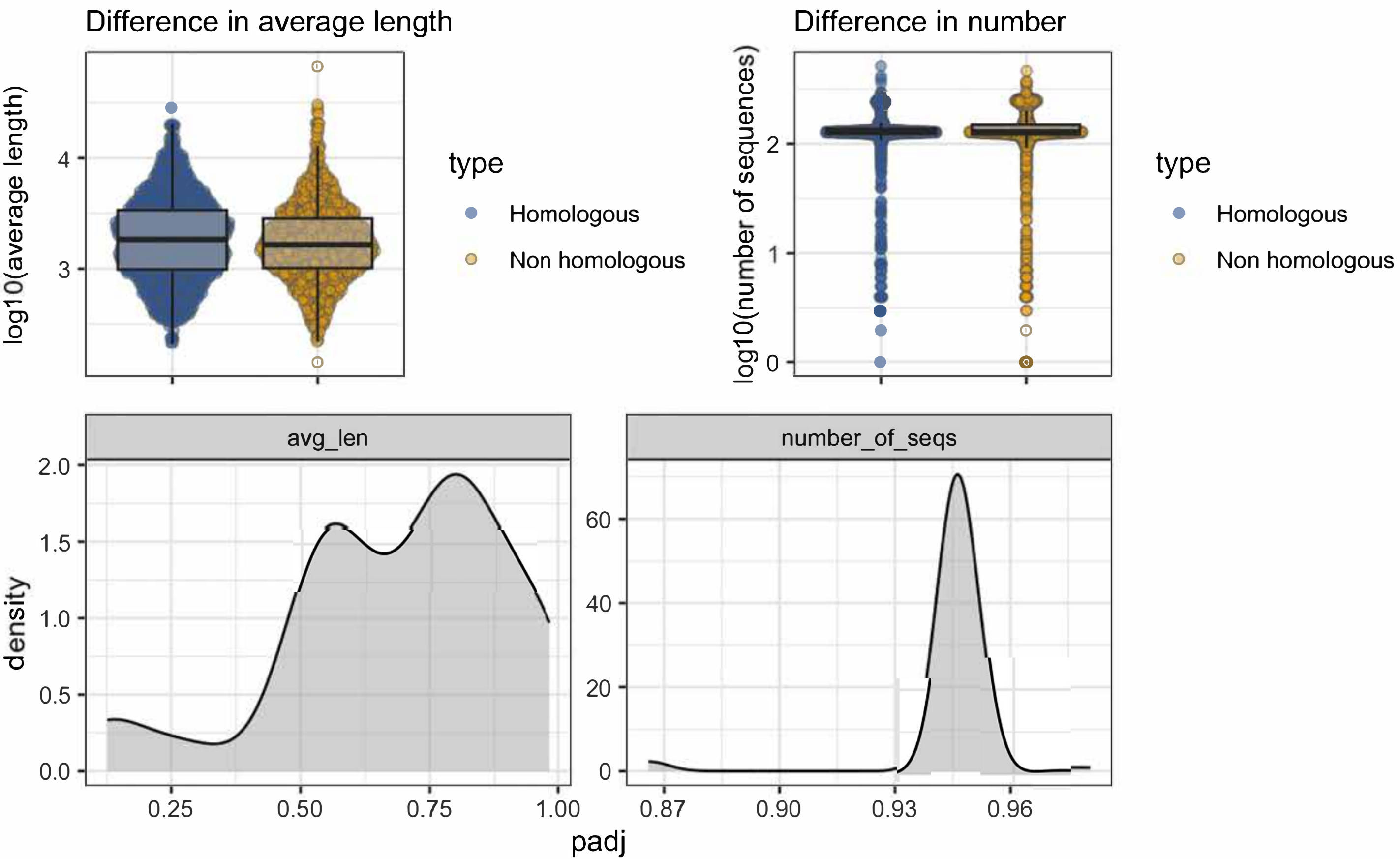

Difference in average length
log10(average length)
4
3
type
Homologous
Non homologous
Difference in number
log10(number of sequences)
2
1
0
type
Homologous
Non homologous
avg_len
number_of_seqs
density
2.0
1.5
1.0
0.5
0.0
60
40
20
0
0.25
0.50
0.75
1.00
0.87
0.90
0.93
0.96
padj

**Fig S7: Checking for possible biases in null set.** We checked any significant difference in average gene length and average number of sequence between genes in the conserved network (homologous set in this plot) and random set of genes not in the conserved network (non- homologous set in this plot). A bootstrapped two-sided Wilcoxon rank-sum test showed no significant differences in copy numbers. The distribution plots bellow shows the disrtibution of Benjamini-Hochberg corrected P-values for the above test. The values are well above the conventional threshold of 0.05.

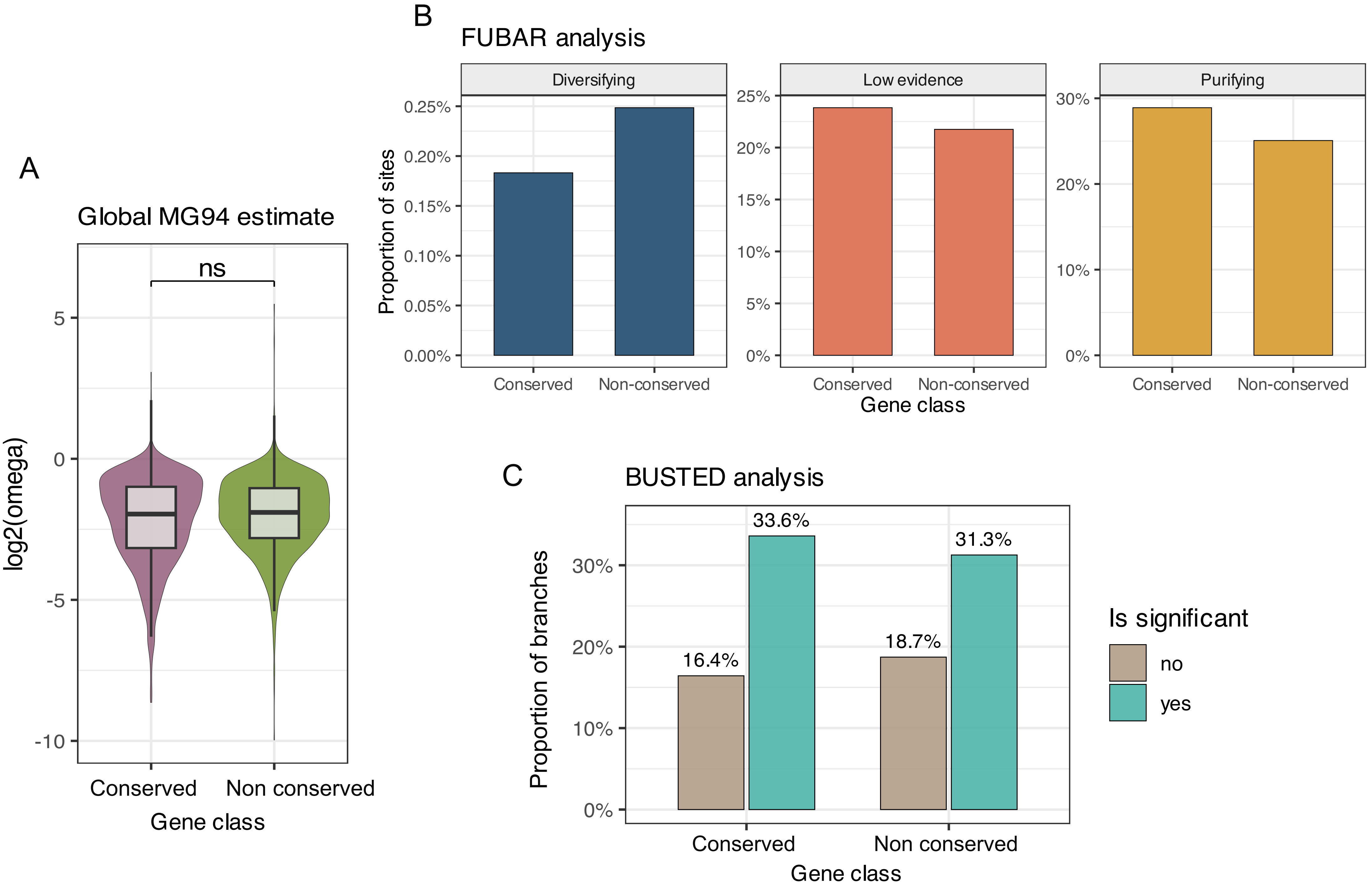
A
Global MG94 estimate
ns
log2(omega)
5
0
-5
-10
Conserved
Non conserved
Gene class
B
FUBAR analysis
Diversifying
Low evidence
Purifying
Proportion of sites
0.25%
0.20%
0.15%
0.10%
0.05%
0.00%
25%
20%
15%
10%
5%
0%
30%
20%
10%
0%
Conserved
Non-conserved
Gene class
C
BUSTED analysis
Proportion of branches
30%
20%
10%
0%
16.4%
33.6%
18.7%
31.3%
Conserved
Non conserved
Gene class
Is significant
no
yes

**Fig S8: Evolutionary analyses of genes in the conserved network.** (A) A global MG94 model showed no difference in dN/dS between genes from the conserved network versus genes not in the conserved network. (B) FUBAR analysis revealed that a higher proportion of sites in genes from the conserved network are under purifying selection suggesting high evolutionary constraint at the coding sequence level. (C) Analysis for episodic diversifying selection using BUSTED showed a higher proportion of branches with evidence of diversifying selection

#### Copy–number variation of high–turnover genes and species traits: all PGLMMs

384 coefficients from 16 modules × 9 traits; 47 have a 95% credible interval excluding 0 (32 after removing associations flagged unreliable).
About 19 would be expected by chance.

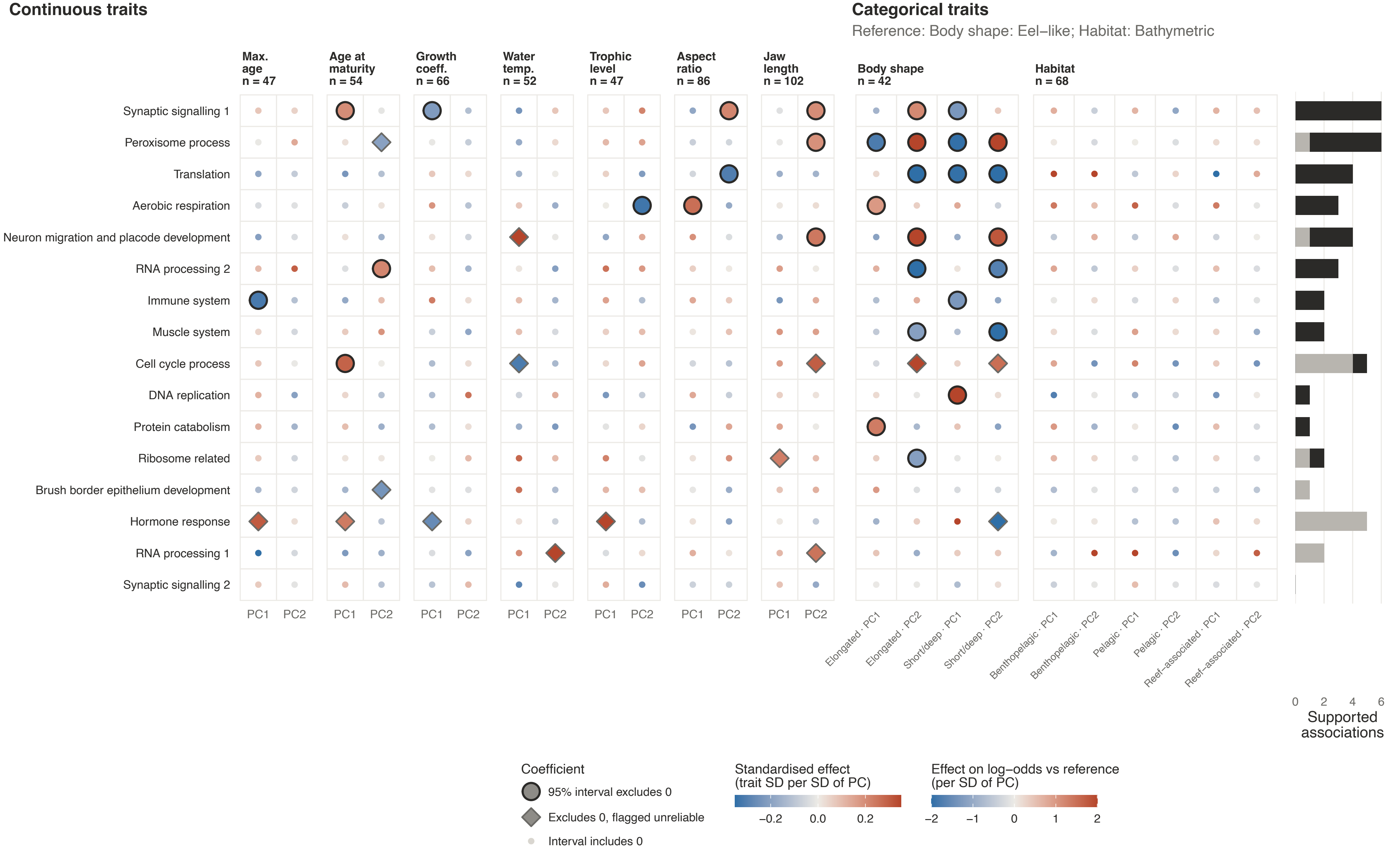


Colour: posterior mean effect, capped at the colour–bar limits. Large ringed circle: 95% interval excludes 0;
diamond: excludes 0 but flagged unreliable in 09_pglmm.Rmd. Right–hand bars: supported associations per module (grey: flagged).
n: species in the model (largest across modules; a few modules have one fewer).

**Fig S9: Overview of results estimates from the phylogenetic generelised linear mixed model.** The y-axis provides the model estimate for each of the respose variables, and the x-axis denoted the predictor variable and levels. The model estimate is only significant if the confidence interval, denoted by the error bars, does not overlap zero. Significant model estimates are denoted by the pink triangles.

**Supplementary text**

**Supplementary text 1:** Gene copy number plot from Fig5B in the main text showed extensive losses in multiple lineages. This extensive loss was surprising. We verified this using the independent orthology estimates from NCBI. Out of the approximately 200 RefSeq teleost genomes, NCBI identified these orthologs in only 65 species for *atp5mea* (https://www.ncbi.nlm.nih.gov/datasets/gene/559659/#orthologs) and 62 species for *atp5meb* (https://www.ncbi.nlm.nih.gov/datasets/gene/767750/#orthologs). Currently, there is no evidence of hidden orthology for these gene. We plan to explore the genomic variation of this gene family in a future study.

**Supplementary text 2: Gene family turnover in cell cycle and synaptic signalling modules are associated with age of maturity and grown coefficient respectively.** We found evidence for a positive relationship between high turnover ortholog groups of the cell cycle process module and age of maturity (Fig S10A). Fish with high age of maturity such as the South Georgia icefish (*Pseudochaenichthys georgianus*; Psgeo), European plaice (*Pleuronectes platessa*; Plpla), and several species of eels (*Anguillia anguilla*; Anang; *Anguilla rostrata*: Anros; *Conger conger*; Cocon) occupy higher positions on PC1, with ortholog groups containing survivin genes (*birc5a*) and TPX2 microtubule nucleation factor (*tpx*) driving most of the variation (Fig S10A). All four species have experienced an expansion in these ortholog groups (Fig S10B). These ortholog groups were typically highly expressed in the gonads across multiple teleost species, as well as mesonephros and head kidney (Fig S10C). Phenotypes of these trait-associated high-turnover genes were largely restricted to the nervous system (Fig S10D).

Another noteworthy relationship was between variation in gene number of high turnover ortholog groups in the synaptic signalling module and von Bertalanffy growth coefficient (Fig S11A). Fast growing fish such as medaka (*Oryzias latipes*), and

Guppy (*Poecilia reticulata* ) occupy lower position along PC1 while slower growing fish such as eels, and blue catfish (*Ictalurus furcatus*) occupy higher positions on PC1. There are multiple high turnover ortholog groups comprising protein tyrosine phosphatase receptors (*ptprr*), and stathmin-like genes (*stmn4*), that are driving variation along PC1 (Fig S11A). However, copy number plots on the phylogeny do not show any distinctive configurations in these ortholog groups (Fig 11B), suggesting a composite effect of copy-number variation across multiple ortholog groups, rather than a distinctive change in any single ortholog group. Expression data from multiple fish species confirms the role of these ortholog groups in the central nervous system with the brain and eye typically being the tissue where these genes have highest expression (Fig 11C). Gene perturbation data from ZFIN showed that disrupting genes making up the synaptic signalling module affects multiple structures across the central nervous (Fig S11D).

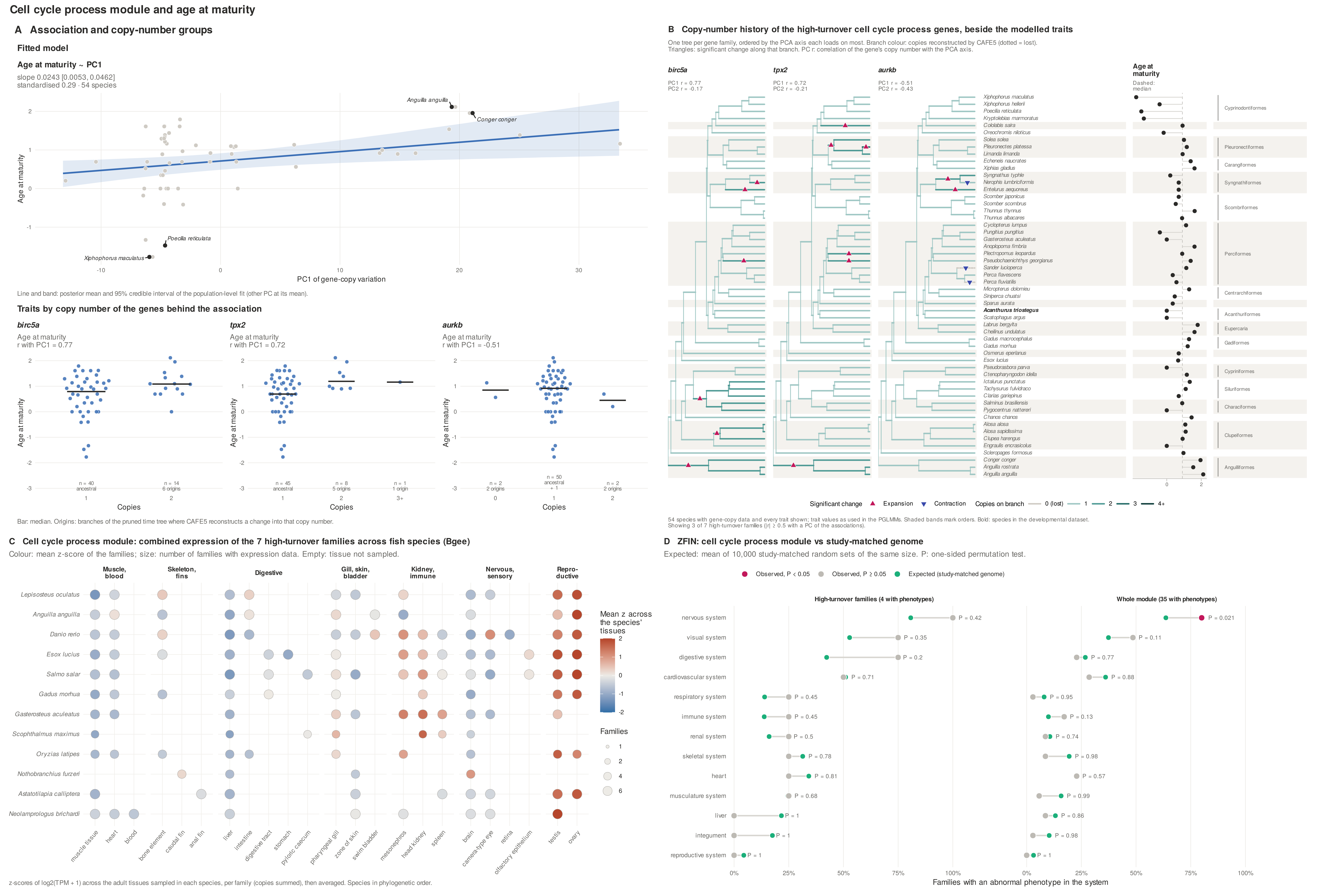

Cell cycle process module and age at maturity
A Association and copy-number groups
Fitted model
Age at maturity ~ PC1
slope 0.0243 [0.0053, 0.0462]
standardised 0.29 · 54 species
Anguilla anguilla
Conger conger
Poecilia reticulata
Xiphophorus maculatus
Age at maturity
PC1 of gene-copy variation
Line and band: posterior mean and 95% credible interval of the population-level fit (other PC at its mean).
Traits by copy number of the genes behind the association
birc5a
Age at maturity
r with PC1 = 0.77
tpx2
Age at maturity
r with PC1 = 0.72
aurkb
Age at maturity
r with PC1 = -0.51
n = 40 ancestral
n = 14 6 origins
n = 45 ancestral
n = 8 5 origins
n = 1 1 origin
n = 2 2 origins
n = 50 ancestral + 1
n = 2 2 origins
Copies
Bar: median. Origins: branches of the pruned time tree where CAFE5 reconstructs a change into that copy number.
B Copy-number history of the high-turnover cell cycle process genes, beside the modelled traits
One tree per gene family, ordered by the PCA axis each loads on most. Branch colour: copies reconstructed by CAFE5 (dotted = lost).
Triangles: significant change along that branch. PC r: correlation of the gene's copy number with the PCA axis.
birc5a
PC1 r = 0.77
PC2 r = -0.17
tpx2
PC1 r = 0.72
PC2 r = -0.21
aurkb
PC1 r = -0.51
PC2 r = -0.43
Age at maturity
Dashed: median
Xiphophorus maculatus
Xiphophorus hellerii
Poecilia reticulata
Kryptolebias marmoratus
Cololabis saira
Oreochromis niloticus
Solea solea
Pleuronectes platessa
Limanda limanda
Echeneis naucrates
Xiphias gladius
Syngnathus typhle
Nerophis lumbriciformis
Entelurus aequoreus
Scomber japonicus
Scomber scombrus
Thunnus thynnus
Thunnus albacares
Cyclopterus lumpus
Pungitius pungitius
Gasterosteus aculeatus
Anoplopoma fimbria
Plectropomus leopardus
Pseudochaenichthys georgianus
Sander lucioperca
Perca flavescens
Perca fluviatilis
Micropterus dolomieu
Siniperca chuatsi
Sparus aurata
Acanthurus triostegus
Scatophagus argus
Labrus bergylta
Cheilinus undulatus
Gadus macrocephalus
Gadus morhua
Osmerus eperlanus
Esox lucius
Pseudorasbora parva
Ctenopharyngodon idella
Ictalurus punctatus
Tachysurus fulvidraco
Clarias gariepinus
Salminus brasiliensis
Pygocentrus nattereri
Chanos chanos
Alosa alosa
Alosa sapidissima
Clupea harengus
Engraulis encrasicolus
Scleropages formosus
Conger conger
Anguilla rostrata
Anguilla anguilla
Cyprinodontiformes
Pleuronectiformes
Carangiformes
Syngnathiformes
Scombriformes
Perciformes
Centrarchiformes
Acanthuriformes
Eupercaria
Gadiformes
Cypriniformes
Siluriformes
Characiformes
Clupeiformes
Anguilliformes
Significant change ▲ Expansion ▼ Contraction
Copies on branch 0 (lost) 1 2 3 4+
54 species with gene-copy data and every trait shown; trait values as used in the PGLMMs. Shaded bands mark orders. Bold: species in the developmental dataset.
Showing 3 of 7 high-turnover families (|r| ≥ 0.5 with a PC of the associations).
C Cell cycle process module: combined expression of the 7 high-turnover families across fish species (Bgee)
Colour: mean z-score of the families; size: number of families with expression data. Empty: tissue not sampled.
Muscle, blood
Skeleton, fins
Digestive
Gill, skin, bladder
Kidney, immune
Nervous, sensory
Repro-ductive
Lepisosteus oculatus
Anguilla anguilla
Danio rerio
Esox lucius
Salmo salar
Gadus morhua
Gasterosteus aculeatus
Scophthalmus maximus
Oryzias latipes
Nothobranchius furzeri
Astatotilapia calliptera
Neolamprologus brichardi
muscle tissue
heart
blood
bone element
caudal fin
anal fin
liver
intestine
digestive tract
stomach
pyloric caecum
pharyngeal gill
zone of skin
swim bladder
mesonephros
head kidney
spleen
brain
camera-type eye
retina
olfactory epithelium
testis
ovary
Mean z across the species' tissues
Families
z-scores of log2(TPM + 1) across the adult tissues sampled in each species, per family (copies summed), then averaged. Species in phylogenetic order.
D ZFIN: cell cycle process module vs study-matched genome
Expected: mean of 10,000 study-matched random sets of the same size. P: one-sided permutation test.
Observed, P < 0.05
Observed, P ≥ 0.05
Expected (study-matched genome)
High-turnover families (4 with phenotypes)
Whole module (35 with phenotypes)
nervous system P = 0.42 P = 0.021
visual system P = 0.35 P = 0.11
digestive system P = 0.2 P = 0.77
cardiovascular system P = 0.71 P = 0.88
respiratory system P = 0.45 P = 0.95
immune system P = 0.45 P = 0.13
renal system P = 0.5 P = 0.74
skeletal system P = 0.78 P = 0.98
heart P = 0.81 P = 0.57
musculature system P = 0.68 P = 0.99
liver P = 1 P = 0.86
integument P = 1 P = 0.98
reproductive system P = 1 P = 1
Families with an abnormal phenotype in the system

**Fig S10: Cell cycle process and age at maturity.** (A) age at maturity increases along PC1 (standardised effect 0.29, 54 species). The genes behind it are *birc5a* and *tpx2* (r = 0.77 and 0.72) and *aurkb* (r = −0.51). Species with an extra *birc5a* or *tpx2* copy appear to mature slightly later. (B) shows 3 of the 7 high-turnover families, those with $|r| \geq 0.5$ on PC1. Most changes are expansions, scattered across the tree. (C) expression is highest in testis and ovary, and in head kidney in some species, which fits its aactivity in proliferating tissue. (D) Weak support for ZFIN the pertubation phenotypes. Four high-turnover families have phenotypes, and none is enriched in any system. The whole module is enriched only for nervous system (P = 0.021).

#### Synaptic signalling 1 module and growth coefficient

##### A Association and copy-number groups

**Fitted model**

**Growth coefficient ~ PC1**

slope -0.0150 [-0.0309, -0.0001]
standardised -0.22 · 66 species

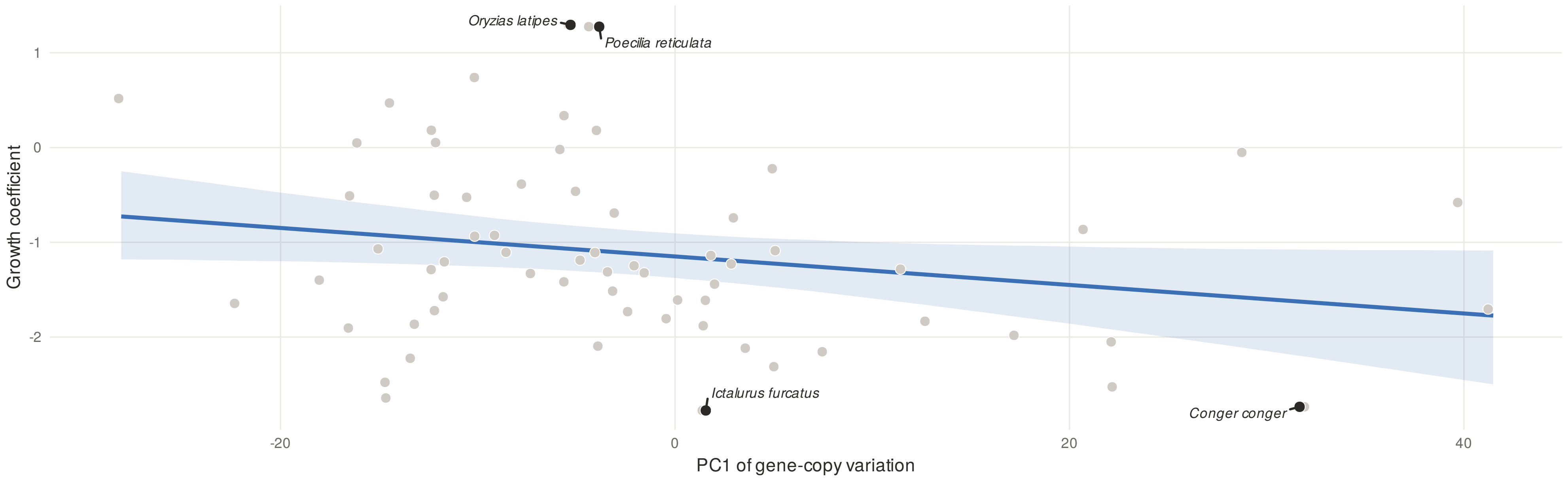


Line and band: posterior mean and 95% credible interval of the population-level fit (other PC at its mean).

**Traits by copy number of the genes behind the association**

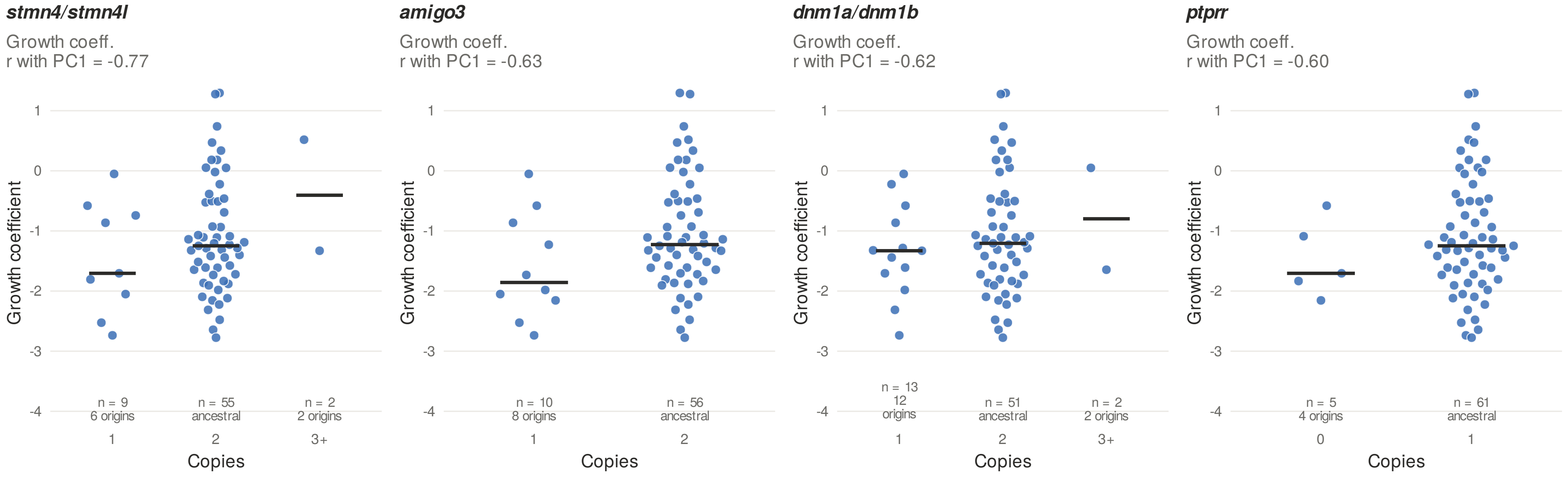


Bar: median. Origins: branches of the pruned time tree where CAFE5 reconstructs a change into that copy number.

##### B Copy-number history of the high-turnover synaptic signalling 1 genes, beside the modelled traits

One tree per gene family, ordered by the PCA axis each loads on most. Branch colour: copies reconstructed by CAFE5 (dotted = lost).
Triangles: significant change along that branch. PC r: correlation of the gene's copy number with the PCA axis.

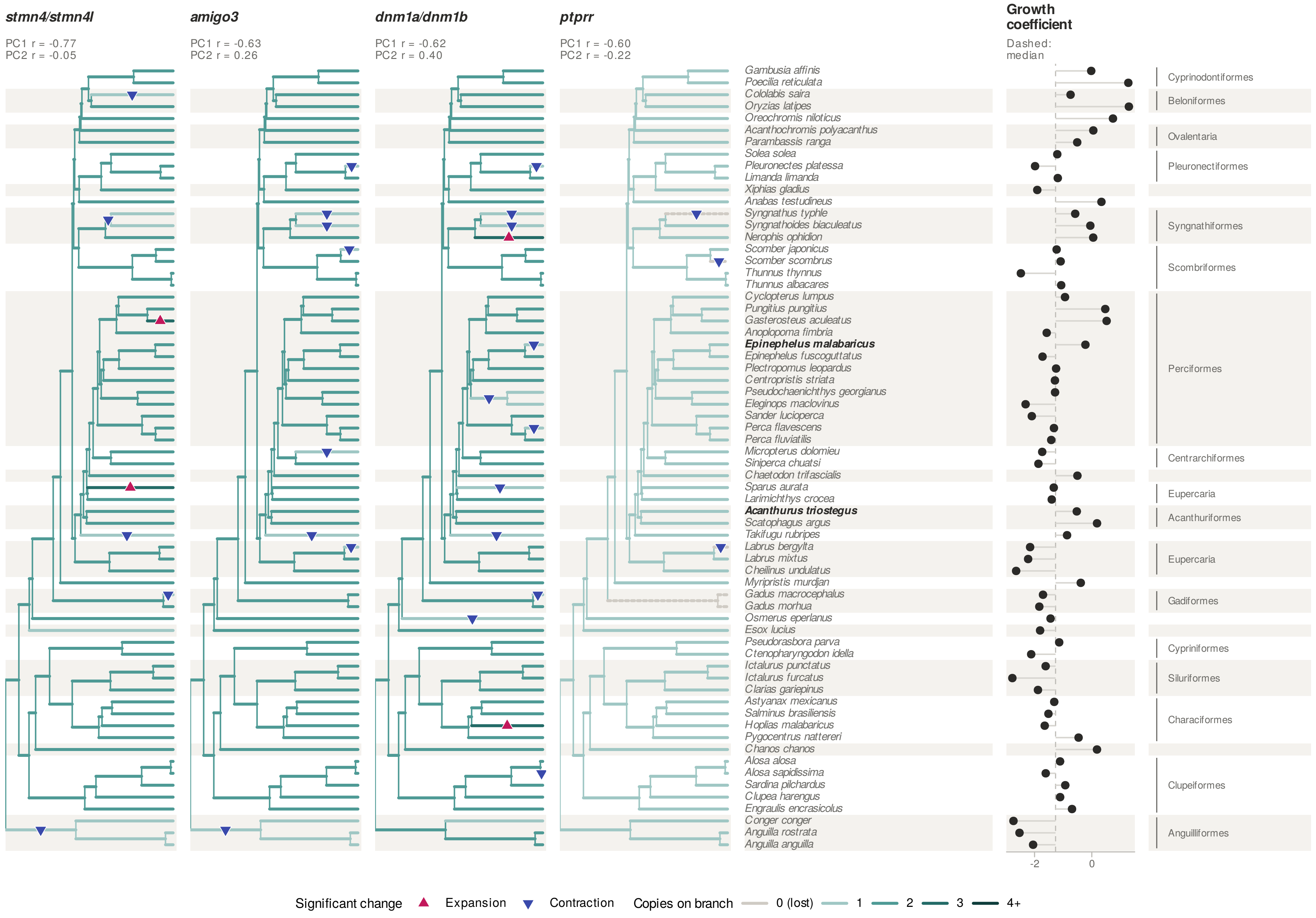


66 species with gene-copy data and every trait shown; trait values as used in the PGLMMs. Shaded bands mark orders. Bold: species in the developmental dataset.
Showing 4 of 11 high-turnover families (|r| ≥ 0.5 with a PC of the associations).

##### C Synaptic signalling 1 module: combined expression of the 11 high-turnover families across fish species (Bgee)

Colour: mean z-score of the families; size: number of families with expression data. Empty: tissue not sampled.

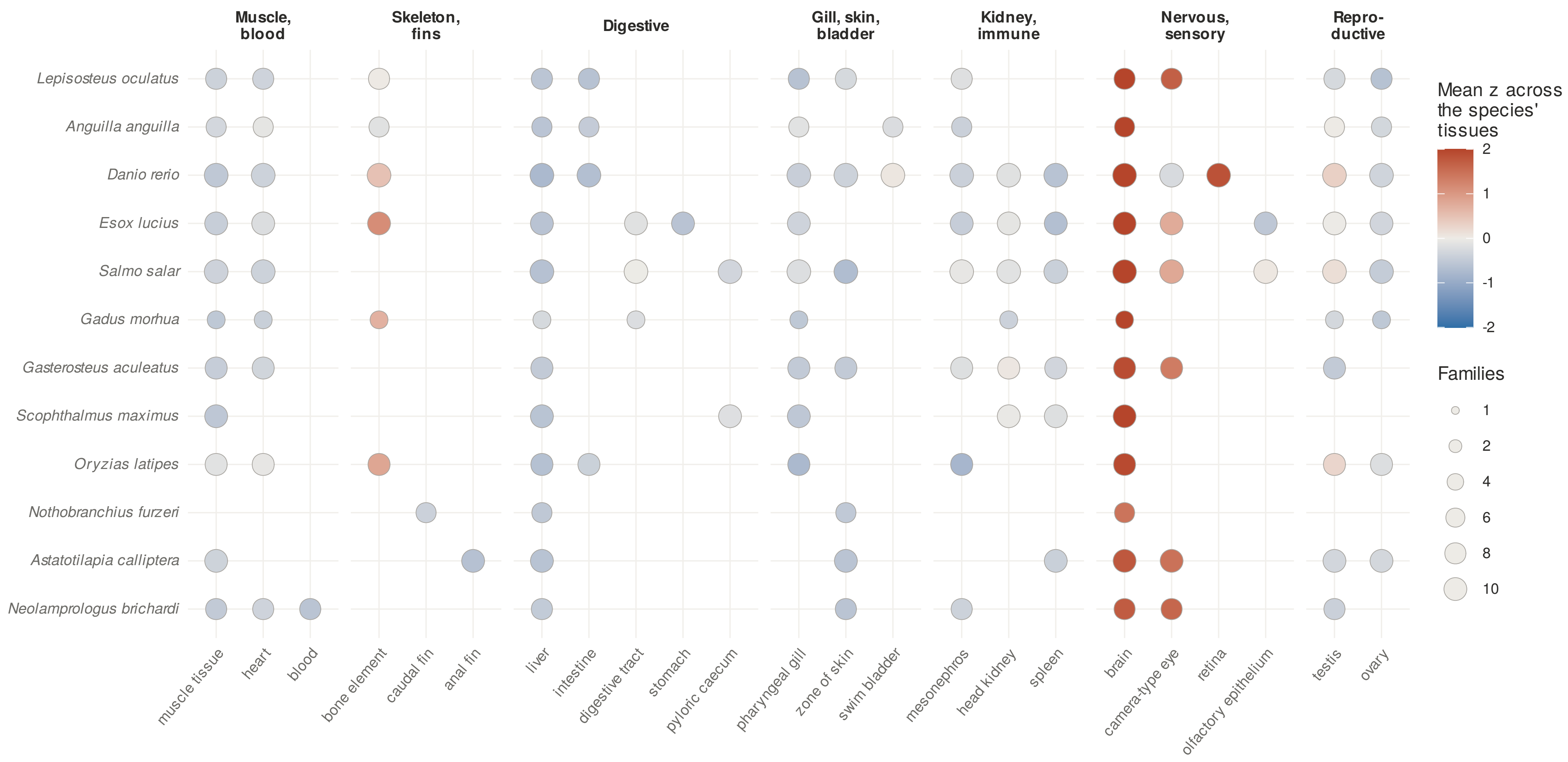


z-scores of log2(TPM + 1) across the adult tissues sampled in each species, per family (copies summed), then averaged. Species in phylogenetic order.

##### D ZFIN: synaptic signalling 1 module vs study-matched genome

Expected: mean of 10,000 study-matched random sets of the same size. P: one-sided permutation test.

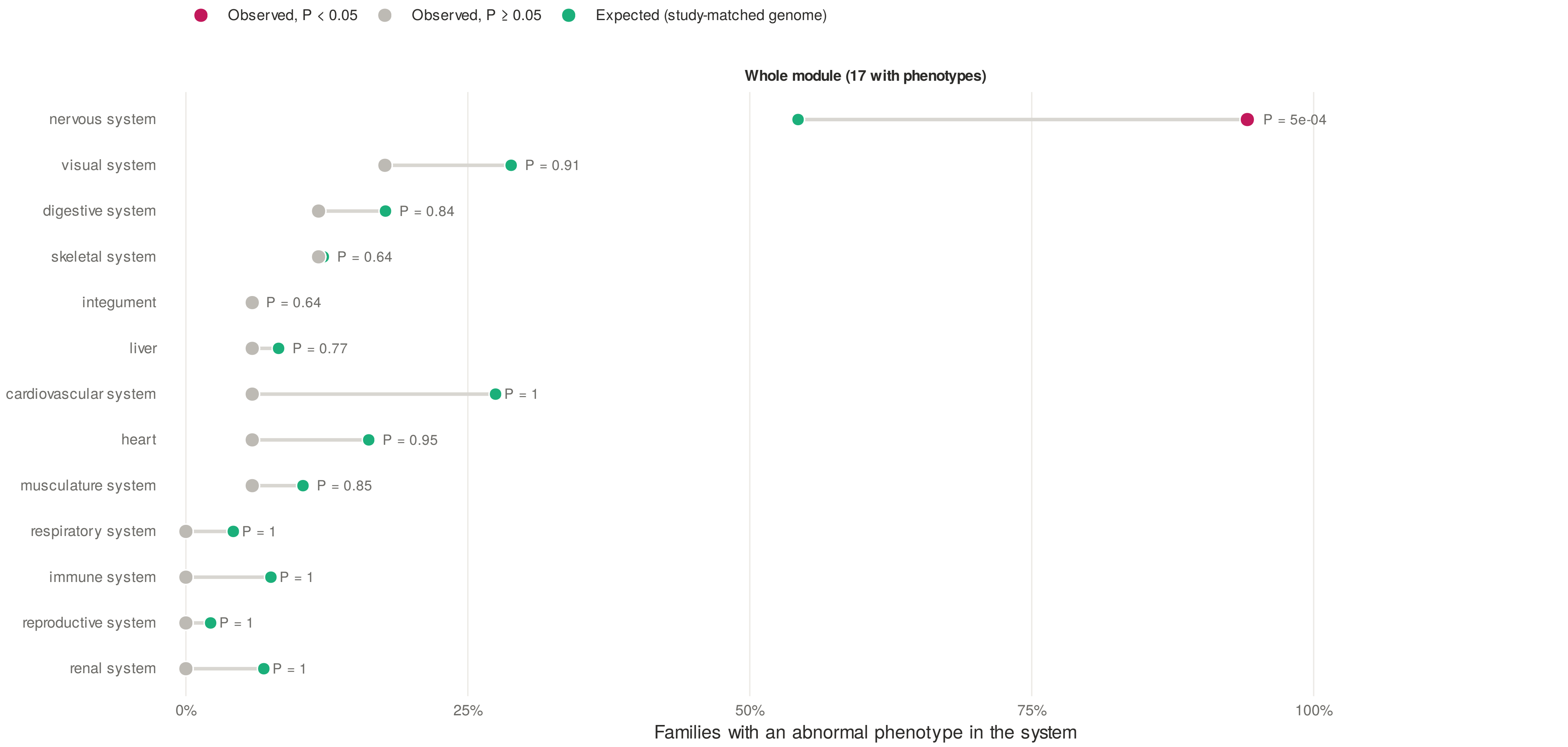


Fewer than 3 high-turnover families have a ZFIN phenotype, so only the whole module is tested.

**Fig S11: Synaptic signalling 1 and growth coefficient. (**A) Growth coefficient decreases along PC1 (standardised effect −0.22, 66 species; the 95% interval only just excludes zero). The genes behind it all load negatively: *stmn4*, *amigo3*, *dnm1a* and *ptprr*. Therefore, species with more copies tend to have a higher growth coefficient. (B) This plot shows 4 of the 11 high-turnover families. Most changes are contractions. (C) Expression across tissues shows a consistent pattern. Brain is the highest tissue in nearly every species, with eye the second highest. (D)ZFIN phenotypes provide strong support. The module as a whole is enriched for nervous-system phenotypes (94% of families vs 54% expected, P = 0.0005), and this survives FDR correction across all modules.

**Supplementary methods**

**Supplementary methods 1: Zebrafish and goby sample collection and sequencing.**

Developmental stages were selected according to the staging scheme of Parichy et al. (2009). For each developmental condition, whole zebrafish larvae were pooled prior to RNA extraction, with the number of individuals per pool adjusted according to specimen size. Specifically, 2–5 individuals were used per replicate, and three biological replicates were collected for each stage. The sampled stages were 6.1 mm, 6.5 mm, 7.1 mm, 7.5 mm, 8.2 mm, 8.5 mm, 8.7 mm, 9.5 mm, 10.3 mm, 13.1 mm, and 16.0 mm standard length. Samples were submiQed to the UMMC Molecular and Genomics Core Facility for RNA isolation, quality control, and bulk RNA sequencing. Total RNA was extracted from whole specimens using TRIzol Reagent in combination with the PureLink RNA Mini Kit (Invitrogen), following the manufacturer's instructions. RNA quality was assessed on the basis of concentration and integrity, including visualization of intact 18S and 28S rRNA bands and an RNA Integrity Score (RIS) greater than 8. Strand-specific mRNA libraries were prepared using the Illumina TruSeq Stranded mRNA Library Prep Kit and sequenced on an Illumina NextSeq 500 using the High Output Kit (150 cycles), generating paired-end 75-bp reads.

For goby, larvae were collected from freshwater streams in northern Okinawa, Japan. Developmental stages were assigned according to the classification of Kondo et al. (2013). Whole larvae were pooled for each sample, with 2–4 individuals per replicate depending on specimen size, and at least three biological replicates were obtained for each developmental group. The seven sampled groups corresponded to 6.0–6.6, 8.0, 8.4, 9.4, 11.4, 16.7, and 25.1 mm standard length. Total RNA was extracted from whole specimens using the Maxwell RSC simplyRNA Tissue Kit (Promega; AS1340). Strand-specific RNA-seq libraries were prepared using the NEBNext Ultra II Directional RNA Library Prep Kit for Illumina. Sequencing was performed at the OIST Sequencing Center on an Illumina NovaSeq 6000 using the SP Reagent Kit v1.0 (300 cycles), generating paired-end 130-bp reads.

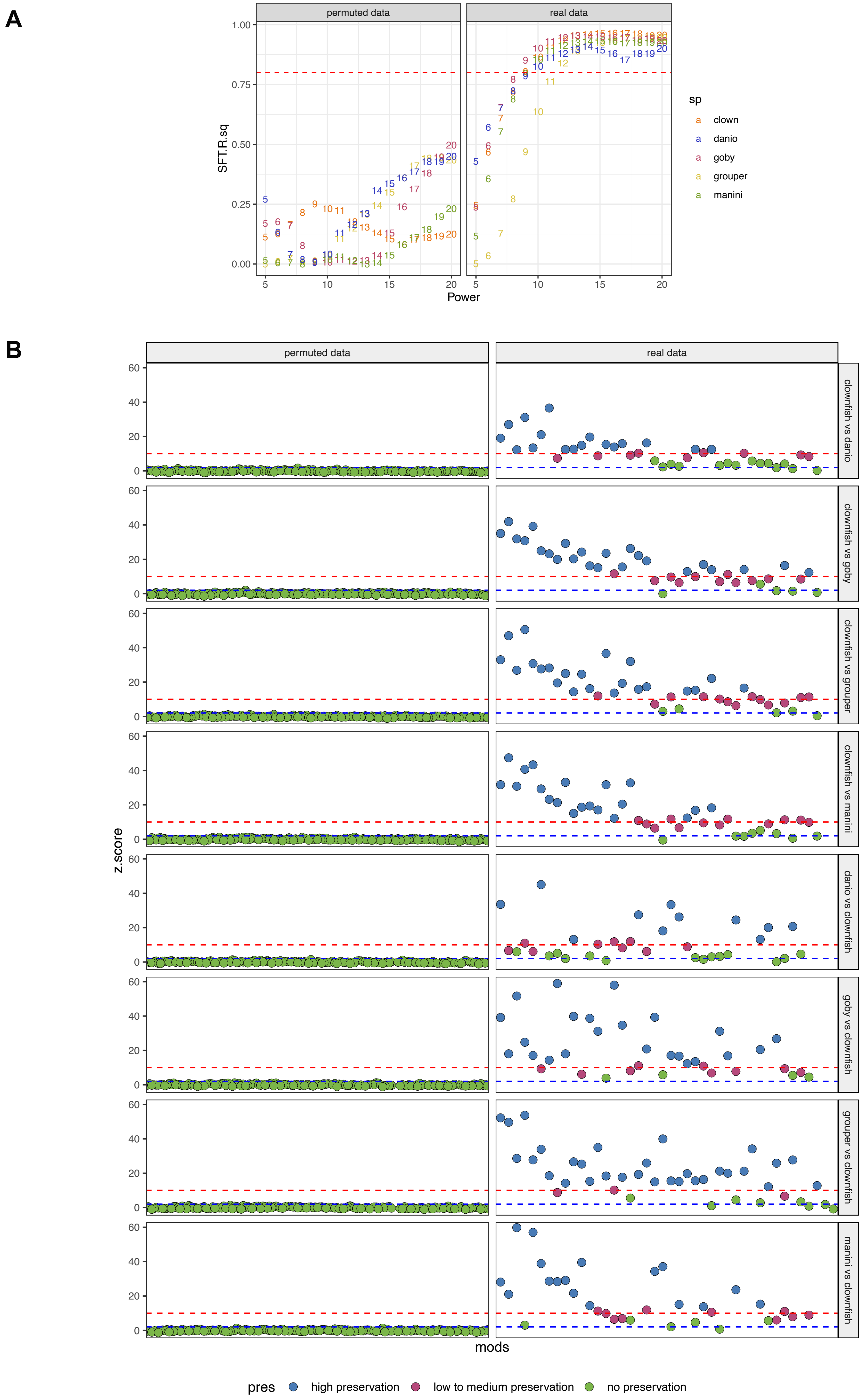
A
permuted data
real data
SFT.R.sq
Power
sp
clown
danio
goby
grouper
manini
B
permuted data
real data
clownfish vs danio
clownfish vs goby
clownfish vs grouper
clownfish vs manini
danio vs clownfish
goby vs clownfish
grouper vs clownfish
manini vs clownfish
z.score
mods
pres
high preservation
low to medium preservation
no preservation

**Supplementary methods 2: Comparisons with a random network.** To ensure that the modules we obtain represent biologically meaningful grouping of genes due to their coordinated expression during post-embryonic development, we constructed random networks using permuted gene expression values and compared the network structures. To make the row-wise permutation for each gene, we randomly shuffled expression values (for each species) across samples. This preserved each gene's expression distribution, mean, and variance, but broke the coordinated expression structure between genes due to development. Using this null we checked for clustering simply due transcriptomic grouping of housekeeping genes. (A) The random networks failed to achieve scale-free topology as denoted by very low correlation (SFT.R.sq) for any power. Scale free topology is an essentail property of biologically relevant networks. The networks using real data achieved very high correlation to scale-free topology structure even with low power. (B) The modules obtained using the permuted data do not show any preservation between species. Therefore, the modules from our original networks are a biologically meaningful representation of coordinated gene activity during post-embryonic development, and not simply transcriptomic grouping of housekeeping genes.

**Supplementary methods 3: T3 and MPI treatments.** The data for the T3 and MPI treatments was obtained from a previous study (30). For the full methods behind the T3 and MPI treatments please refer to (30). In brief, A. ocellaris larvae were exposed to triiodothyronine (T3; 48h), goitrogen mixture MPI (methimazole and potassium perchlorate; 19days), and DMSO as control. Iopanoic acid was included with T3 which was required to inhibit deiodinase activity. Larvae were maintained in groups of 10 in 800-ml beakers at 27°C and fed rotifers three times daily (10 rotifers $ml^{-1}$) and Artemia nauplii once daily. Water (100 ml) was replaced daily, and treatment compounds were replenished to maintain constant concentrations. Larvae were treated for 48h with DMSO and T3 ($10^{-6}$ M) and sampled for transcriptomic analysis. Larvae were treated for 19 days with DMSO and MPI (diluted 1:1000) and sampled for transcriptome analysis. Treatment concentrations were selected based on previous experiments investigating fish metamorphosis (27,43).
